# Antibody avidity and multi-specificity combined to confer protection against SARS-CoV-2 and resilience against viral escape

**DOI:** 10.1101/2022.10.23.513379

**Authors:** Clare Burn Aschner, Krithika Muthuraman, Iga Kucharska, Hong Cui, Katherine Prieto, Manoj S. Nair, Maple Wang, Yaoxing Huang, Natasha Christie-Holmes, Betty Poon, Jessica Lam, Azmiri Sultana, Robert Kozak, Samira Mubareka, John L. Rubinstein, Edurne Rujas, Bebhinn Treanor, David D. Ho, Arif Jetha, Jean-Philippe Julien

## Abstract

SARS-CoV-2, the causative agent of COVID-19, has been responsible for a global pandemic. Monoclonal antibodies have been used as antiviral therapeutics, but have been limited in efficacy by viral sequence variability in emerging variants of concern (VOCs), and in deployment by the need for high doses. In this study, we leverage the MULTI-specific, multi-Affinity antiBODY (Multabody, MB) platform, derived from the human apoferritin protomer, to drive the multimerization of antibody fragments and generate exceptionally potent and broad SARS-CoV-2 neutralizers. CryoEM revealed a high degree of homogeneity for the core of these engineered antibody-like molecules at 2.1 Å resolution. We demonstrate that neutralization potency improvements of the MB over corresponding IgGs translates into superior *in vivo* protection: in the SARS-CoV-2 mouse challenge model, comparable *in vivo* protection was achieved for the MB delivered at 30x lower dose compared to the corresponding IgGs. Furthermore, we show how MBs potently neutralize SARS-CoV-2 VOCs by leveraging augmented avidity, even when corresponding IgGs lose their ability to neutralize potently. Multiple mAb specificities could also be combined into a single MB molecule to expand the neutralization breadth beyond SARS-CoV-2 to other sarbecoviruses. Our work demonstrates how avidity and multi-specificity combined can be leveraged to confer protection and resilience against viral diversity that exceeds that of traditional monoclonal antibody therapies.

## Introduction

Emerging infectious agents, including viruses such as SARS-CoV-2, present enormous challenges to global public health through the lack of pre-existing immunity in the population. Despite the availability of vaccines against SARS-CoV-2 disease (COVID-19), global vaccine coverage remains low, with only 19.9% of people in low-income countries having received at least one dose^1^. Relatively short-lived vaccine-mediated protection, coupled with the emergence of new viral variants, further highlights the necessity for effective prophylactic and treatment options^2–10^. Monoclonal antibodies (mAbs), which have been efficacious in the treatment of infectious diseases including respiratory syncytial virus (RSV)^11^ and Ebola virus^12^, present a promising option. Some mAbs, including Bamlanivimab and Etesevimab delivered together^13^, and the REGEN-COV cocktail of Casirivimab and Imdevimab^14^, received US Food and Drug Administration (FDA) authorization to treat COVID-19, but have struggled to overcome viral diversity, and are limited by the requirement for high doses and intravenous administration^15^. Both combinations had their authorization revoked following the emergence of the Omicron BA.1 variant of concern (VOC), which has 37 mutations within the spike domain and 15 mutations within the receptor binding domain (RBD), the target of most clinical antibodies against SARS-CoV-2^16^. To date, only one mAb, Bebtelovimab, retains adequate *in vitro* activity against circulating Omicron subvariants and is FDA-authorized for use against SARS-CoV-2^17, 18^. Authorization has also been updated to allow for an increased dose of a cocktail of Tixagevimab and Cilgavimab, which is expected to maintain activity against subvariants despite a loss of potency at the original dose^19, 20^. Despite these limited authorizations, a number of additional antibodies targeting SARS-CoV-2 spike epitopes, including some with broad neutralization against sarbecoviruses, have been identified^21–28^. However, such increases in mAb breadth are often associated with a reduction in potency^25^, highlighting the necessity of identifying therapeutics that combine potency and breadth.

Increasing antibody valency is a promising approach to enhance apparent binding affinity^29–31^, potentially lowering therapeutic dose, improving breadth, and allowing administration through alternative routes, such as subcutaneous or intramuscular delivery. Aiming to exploit avidity to enhance antibody functional responses, a wide range of antibody engineering strategies have been described^32^. Among those, biologics assembled based on IgM^33^, synthetic nanocages^34^ and Minibinder^35^ formats have demonstrated superior neutralization properties against SARS-CoV-2 compared to conventional mAb formats. Additionally, the avid molecules GEN3009^35^, INBRX-106(Inhibrx)^36^ and IGM-8444^37^ are being tested in Phase I/II clinical trials for the treatment of hematological and solid tumors, highlighting the clinical benefit of multivalent antibody-presenting formats. Following a similar principle, but using the human light-chain apoferritin protomer to drive oligomerization of antibody fragments, we developed a platform called the Multabody (MB) to increase neutralization potency of antibodies targeting SARS-CoV-2^38^ and HIV-1^39^. Using this platform, enhanced affinity can be coupled with multi-specificity – the inclusion of several antibody fragments recognizing different epitopes – to result in antigen recognition that is more resistant to viral mutations^38, 39^. This is particularly relevant in light of immune pressure driving the continued emergence of new variants of SARS-CoV-2, including those against which existing vaccines and drugs are less efficacious^40–42^.

Here, we explored whether the increased *in vitro* SARS-CoV-2 neutralization previously reported for the Multabody could translate to *in vivo* protection at low doses. In addition, we assessed whether MBs could rescue the loss in neutralization potency observed for conventional mAbs against different VOCs and expand breadth beyond SARS-CoV-2. Our data provide proof-of-concept that the MB is a tractable platform that harnesses avidity to provide gains in both the *in vitro* and *in vivo* potency and breadth of antibody-based molecules against SARS-CoV-2 and beyond.

## Results

### Assembly of the tri-specific 298-52-80 Multabody defined at atomic resolution

We have previously reported the generation of tri-specific MB molecules using an engineered apoferritin split design (Fig. 1A)^38, 39^, whereby the human apoferritin protomer was split into two halves based on its four-helical bundle fold: the two N-terminal α helices (N-Ferr) and the two C-terminal α helices (C-Ferr). The genetic fusion and transfection into mammalian cell expression systems of a single chain (sc) Fab or a scFc at the N terminus of each apoferritin half and full apoferritin resulted in the secretion of self-assembled, oligomeric molecules capable of ultrapotent neutralization. Specifically, a tri-specific MB incorporating antibody specificities 298, 52 and 80 increased neutralization potency by ∼1000-fold compared to the corresponding IgG cocktail^38^. This tri-specific MB was described to have antibody-like biochemical properties as assessed in biophysical characterizations after purification and under accelerated thermal stress^38^. To obtain molecular insights into the assembly of the Multabody design, we next characterized this tri-specific MB by cryo-electron microscopy (cryoEM) (Fig. S1).

**Figure 1.**
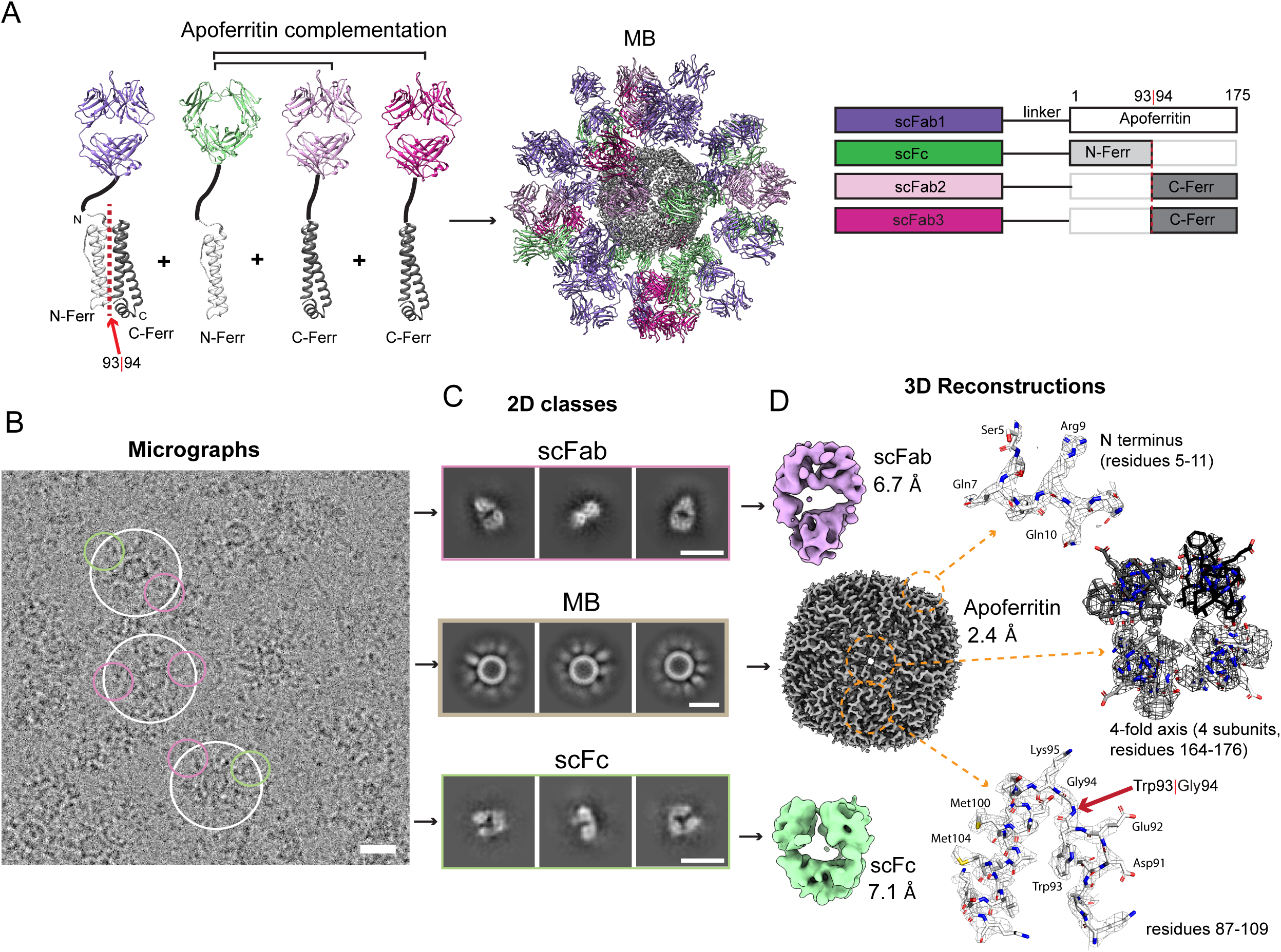
cryoEM characterization of tri-specific MB. (A) Schematic representation of the elements that drive assembly of the tri-specific MB (298-52-80). The red arrow indicates the split site between the first half (N-Ferr) and the second half (C-Ferr) of the human apoferritin light chain. (B) Representative cryoEM micrograph of the tri-specific MB with white and blue circles highlighting the whole particles and scFab/scFc fragments, respectively. (C) Representative 2D classes of scFab (top panels), tri-specific MB (middle panels), and Fc domains (bottom panels). (D) Left panels – cryoEM maps of scFab at 6.7 Å resolution (top), apoferritin nanocage scaffold at 2.4 Å resolution (middle) and scFc at 7.1 Å resolution (bottom). Right panels – fitting of the human apoferritin light chain model (PDB ID:6WX6) into the 2.4 Å map, focusing on features such as the N terminus of the apoferritin scaffold that shows weak density beyond Ser5 due to the flexibility of the linker (top), the four-fold axis formed by four adjacent subunits (middle), and residues 87-109 of the human apoferritin light chain (bottom) where the red arrow points to the split site, between residues Trp93 and Gly94 in some subunits. All cryoEM maps in this figure were refined with no symmetry applied. Scale bars are 10 nm.

Analysis of cryoEM micrographs revealed the formation of highly decorated and homogeneous nanocage-like particles (Fig. 1B). Consistent with the presence of flexible (GGS)_x_ linkers connecting the scFab and scFc components to the apoferritin scaffold, the density of these antibody fragments is poorly resolved in 2D classes (Fig. 1C) and 3D reconstruction of the tri-specific MB (Fig. 1D). However, manual picking of the scFab and scFc particles, followed by template-based particle picking, and subsequent refinement of these molecules confirmed the proper assembly of Fab and Fc components on the MB to ∼7 Å resolution (Fig. 1B-D, Figs. S1 and S2).

3D reconstructions of the apoferritin scaffold of the MB reached 2.4 Å and 2.1 Å resolution, respectively, when no symmetry (C1; Fig. 1D, Fig. S3A-D) or octahedral symmetry (O; Fig. S3E-H) was applied. The apoferritin scaffold in the tri-specific MB is virtually identical to that of the human apoferritin light chain (PDB ID: 6WX6) with measured cross-correlation (cc) coefficients between maps of 0.97 (C1) and 0.92 (O). The N and C termini of the core MB scaffold are similarly disposed in 3- and 4-fold symmetry axes as in the native human apoferritin light chain (Fig. 1D), indicating minimal impact for scFab and scFc genetic fusions. Moreover, the cryoEM maps showed no evidence of deviation from the apoferritin fold for structural elements at the split design site (between residues Trp93 and Gly94; Fig. 1D, bottom right panel). In summary, our cryoEM analysis of the tri-specific 298-52-80 MB provided atomic-level details demonstrating that the MB, built on the apoferritin split design scaffold, adopted its intended structural disposition.

### Neutralization potency correlates with in vivo protection from SARS-CoV-2

Next, we investigated the ability of the tri-specific 298-52-80 MB to confer *in vivo* protection against SARS-CoV-2. To specifically assess the effect of neutralization potency on *in vivo* protection from lethal SARS-CoV-2 challenge, we generated a tri-specific 298-52-80 MB and the corresponding IgG cocktail with an IgG4 Fc containing mutations (S228P, F234A, L235A, G237A, P238S)^43^ to ablate binding to Fcγ receptors, hereafter referred to as MB* and IgG4*, respectively. As expected, replacement of the Fc subtype from IgG1 to IgG4* did not affect the neutralization potency of the IgG or the MB, and, as previously reported^38^. The tri-specific MB* achieved an IC_50_ value of 0.0002 µg/mL, approximately 1000-fold more potent than its corresponding cocktail IgG (Fig. 2A). Binding kinetics studies revealed that both the tri-specific MB* and the IgG4* antibody cocktail displayed pH-dependent binding to mouse and human FcRn (Fig. 2B-C), and no binding to human and mouse Fcγ receptors (FcγR). This was in contrast to the FcγR binding observed for the corresponding IgG1 antibody cocktail control (Fig. 2C, Fig. S4A-B). Antibody-dependent cell-mediated phagocytosis (ADCP) experiments using fluorescently labeled beads coated with SARS-CoV-2 Spike protein further confirmed the inability of the tri-specific MB* and the IgG4* cocktail to engage Fc receptors, while the IgG1 antibody cocktail showed substantial uptake of SARS-CoV-2 Spike-coated beads (Fig. 2D).

**Figure 2.**
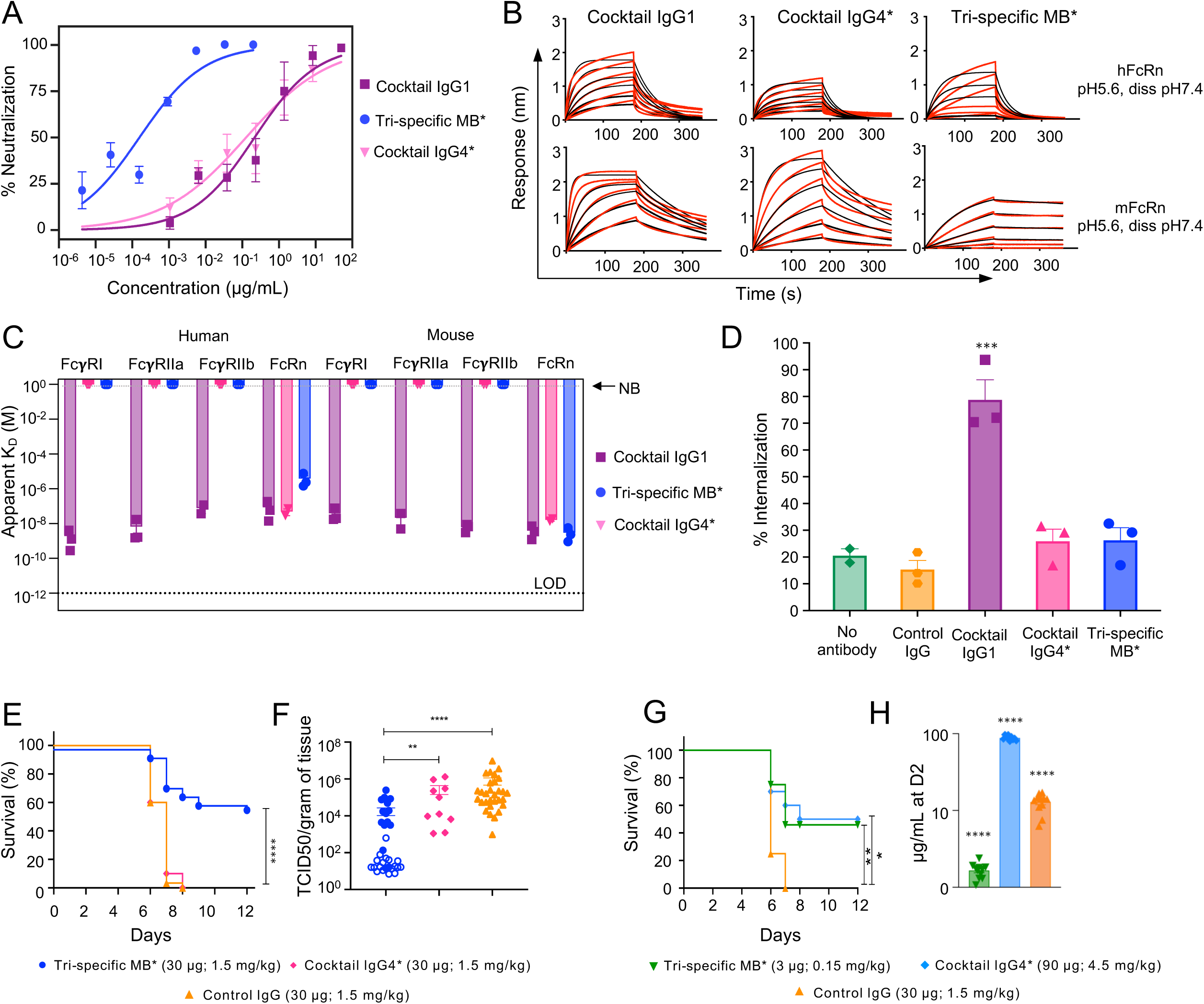
Multabodies protect against SARS-CoV-2 challenge. (A) SARS-CoV-2 PsV neutralization potency of the tri-specific (289-52-80) MB* and the corresponding IgG1 and IgG4 cocktails. * indicates the use of an IgG4* Fc bearing the specified mutations (S228P, F234A, L235A, G237A, P238S) to ablate Fcγ receptor binding. (B) Sensograms of samples shown in (A) binding to human and mouse FcRn (association at pH 5.6, dissociation at pH 7.4). (C) Binding (apparent K_D_) of cocktail IgG1, cocktail IgG4* and tri-specific MB* particles to human (FcγR I, IIa, IIb and FcRn) and mouse (FcγR I, IIb, IV and FcRn) receptors. NB and LOD denote no-binding and limit-of-detection, respectively. (D) ADCP determined as the percentage of THP-1 cells with internalized SARS-CoV-2 Spike-coated fluorescent microspheres. (E-J) Mice expressing hACE2 and hFcRn were dosed (30 µg; 1.5 mg/kg) as indicated and challenged with 1 x 10^5^ PFU / mouse SARS-CoV-2 intranasally. Survival (E) was monitored for 12 days following challenge. (F) Lung viral titers at the end of the experiment (open symbols) or at the time of death in the animals that succumbed (closed symbols) measured by viral outgrowth assay. TCID50 per gram of tissue is shown. Survival (G) and serum IgG or MB concentrations at day 2 (H) following administration of low dose tri-specific MB* (3 µg; 0.15 mg/kg) compared to high-dose (90 µg; 4.5 mg/kg) cocktail IgG4*. Mean values ± SD (A,C-D) and representative data (B) for at least three independent experiments are shown. (D) *** indicates significance compared to no antibody control (p<0.001) by ANOVA. (E-F) n = 33 for tri-specific MB*, n = 30 for control IgG, n = 10 for cocktail IgG4*, from 2-6 independent experiments. (G-H) n = 24 for tri-specific MB*, n = 10 for cocktail IgG4* mix and control IgG, from 2-5 independent experiments. For (E) **** p<0.0001, **p<0.01, *p<0.05 by Gehan Breslow Wilcoxon test. (F) ****p<0.0001, **p<0.01, Kruskall Wallis test and (H) ****p<0.001 compared to all other groups by ANOVA.

To assess whether the increased neutralization potency achieved with the MB resulted in improved *in vivo* protection against SARS-CoV-2, hACE2 and hFcRn double transgenic mice were treated with 30 µg (1.5 mg/kg) of the FcγR-binding deficient IgG4* and MB* molecules and challenged intranasally with a high dose (1 x 10^5^ TCID50) of SARS-CoV-2^44^. The tri-specific MB* provided significantly better protection (60% survival) compared to the IgG4* cocktail, with all cocktail-recipient animals succumbing to the challenge at D6-7 (Fig. 2E). Improved protection was associated with fewer clinical signs of disease throughout the course of infection (Fig. S4D), reduced weight loss and weight rebound following challenge (Fig. S4E), and significantly lower lung viral titers, particularly in animals that survived the challenge (Fig. 2F). Infection was confirmed by qPCR using oropharyngeal swabs collected at D-1 (before challenge) and D2 following challenge (Fig. S4F). In subsequent studies, we found that comparable *in vivo* protection was achieved when the tri-specific MB* was delivered at 3 µg (0.15 mg/kg) and the IgG4* cocktail at 90 μg (4.5 mg/kg) (Fig. 2G). The difference in dose can be observed in circulating serum concentrations of administered molecule at D2 post challenge (Fig. 2H). This data not only provides the first evidence of *in vivo* protection from lethal challenge mediated by the MB, but also illustrates that the increased neutralization potency conferred by the tri-specific MB* format provides enhanced protection against SARS-CoV-2 challenge compared to a corresponding IgG mixture.

### MBs display enhanced potency and breadth against SARS-CoV-2 VOCs

Since the early discovery of mAbs 298, 52, and 80 in 2020, extensive research efforts worldwide have focused on the identification of potent antibodies against SARS-CoV-2^45^. In this context, 2-7^46^, 2-36^24^, 2-38^46^, 10-40^47^ and 11-11^47^ have emerged as RBD-binding mAbs displaying both potency and breadth against SARS-CoV-2 and its variants. To assess whether the neutralization properties of these mAbs could be further improved, we expressed mono-specific MBs and evaluated their neutralization potency and breadth compared to their corresponding mAbs against SARS-CoV-2 wildtype (WT) and five VOCs (Alpha, Beta, Gamma, Delta, and Omicron BA.1). Five of the eight mAbs (52, 80, 2-36, 11-11 and 10-40) showed 100% breadth with an IC_50_ cut-off value of 5 µg/mL. However, when using an IC_50_ cut-off value of 0.01 µg/mL to resemble the potency of REGEN-COV^48^, only two mAbs (11-11 and 10-40) showed neutralization against two of the five VOCs tested (Fig. 3A, Fig. S5). In contrast, when displayed as mono-specific MBs, three specificities (2-7, 80 and 52) reached 100% neutralization breadth using an IC_50_ cut-off value of 0.01 µg/mL. The remaining MBs lose potency against Omicron BA.1, but, apart from 298 and 2-38 MBs, still neutralize with an IC_50_ below 0.3 µg/mL, which represents the pseudovirus (PsV) neutralization potency of sotrovimab ^49^ against WT SARS-CoV-2 (Fig. 3B-C, Fig. S5). The superior ability of these molecules to overcome viral sequence diversity is likely due to their enhanced potency against WT SARS-CoV-2, which ranges from 0.005 to 0.0002 µg/mL (Fig. 3B).

**Figure 3.**
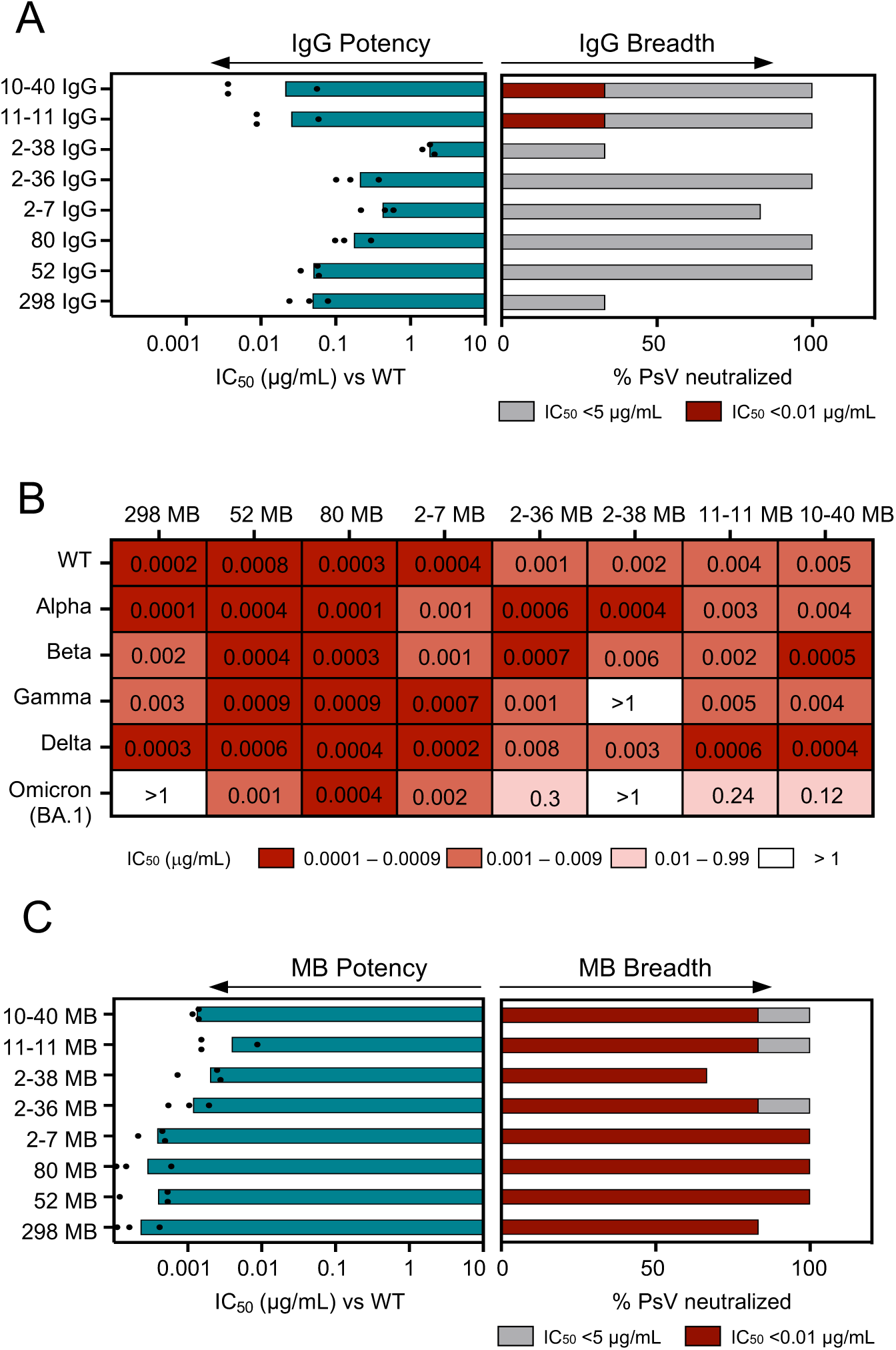
Multabodies are ultrapotent and broad SARS-CoV-2 neutralizers. (A) IgG neutralization potency (teal bars) and breadth against a six PsV panel using a cut-off IC_50_ value of 5 µg/mL (grey bars) or 0.01 µg/mL (red bars). (B) Heat map showing the neutralization potency of mono-specific MBs displaying Fab specificities from (A) against each PsV variant in the panel. Individual IC_50_ values are displayed. (C) Neutralization potency and breadth of mono-specific MBs as in (A). PsV panel: WT, Alpha, Beta, Gamma, Delta, and Omicron BA.1. Data from three biological replicates are shown, bars indicate the mean.

### Molecular basis of Fab 80 binding to SARS-CoV-2 RBD

Next, we sought to understand the molecular basis of binding of mAb 80, as its structure had remained elusive. We solved the crystal structure of 80 Fab in complex with RBD at 3.1 Å resolution (Table S1, Fig. S6). Epitope recognition is mediated by 20 residues that form the interface with the RBD, 14 of which are involved in ACE2 binding (Table S2). This illustrates how mAb 80 inhibits SARS-CoV-2 infection through receptor blockade, preventing the interaction of ACE2 with the receptor binding motif (Fig. 4A). The heavy chain of mAb 80 is primarily responsible for the interaction with RBD, contributing ten of the eleven hydrogen bonds found in the binding interface (Fig. S7A-B, Table S2). Additionally, interaction of F54 of the antibody heavy chain with Y489 from the RBD results in the formation of a new triple pi-stacking within the RBD structure, between residues Y473, F456 and Y421 (Fig. S7C).

**Figure 4.**
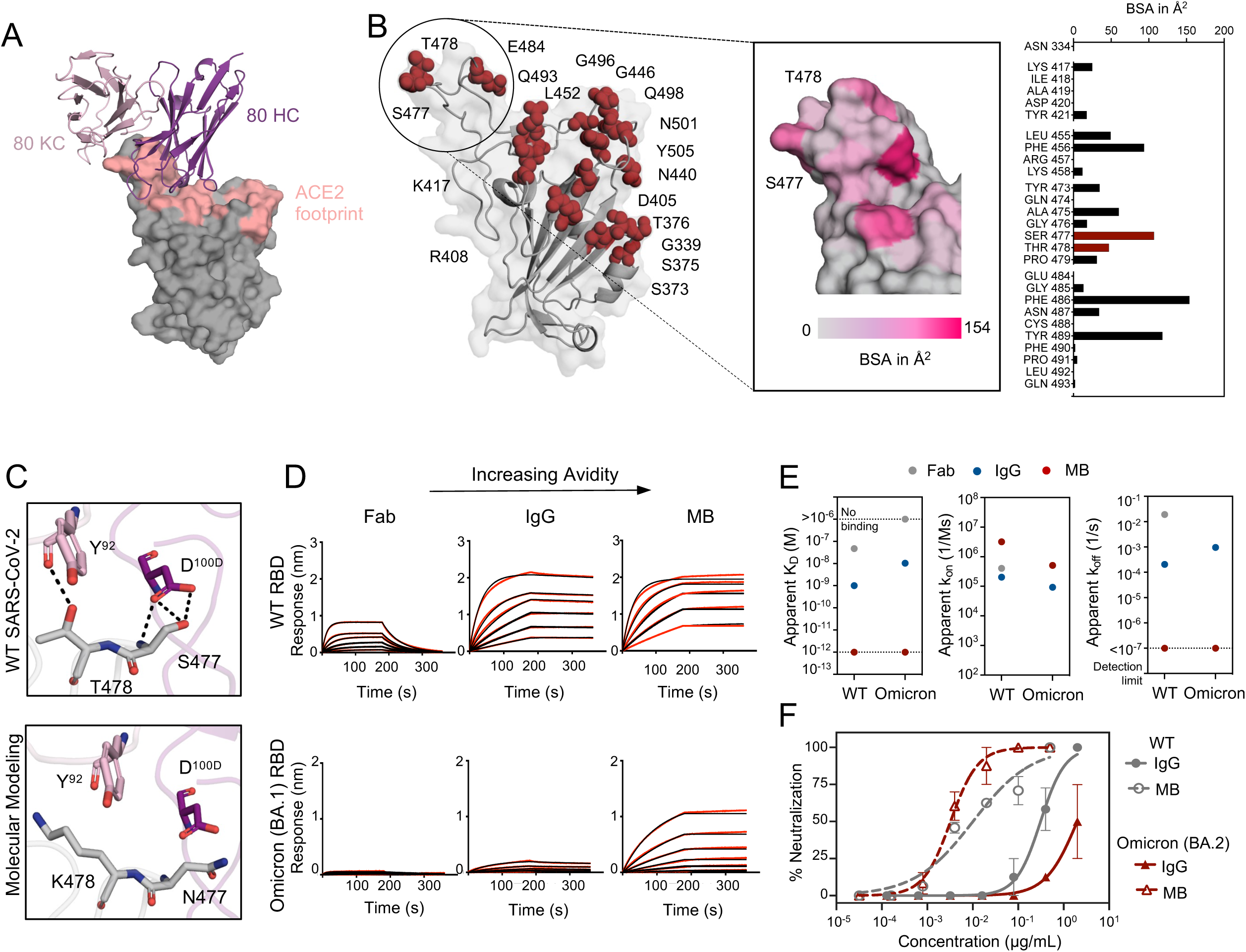
Molecular basis for the resilient SARS-CoV-2 neutralization of 80 MB. (A) Three-dimensional structure of the 80 Fab-RBD complex. The heavy and light chains of 80 Fab are colored in dark and light purple, respectively. RBD is shown as surface representation (grey) with the footprint of ACE2 depicted in salmon. (B) Secondary structure cartoon representation of RBD, with residues mutated in the different variants of concern highlighted as red spheres. Inset: close-up view of the RBD area recognized by 80. Critical residues for binding are colored in pink according to their BSA. Mutated residues in the VOCs are indicated and shown in red in the BSA plot. (C) Molecular modeling displays the possible conformation adopted by the side chains of the mutated residues T478K and S477N upon 80 binding. Hydrogen bonds are shown as dashed black lines. (D) Sensograms of 80 Fab, IgG and MB binding to WT and Omicron BA.1 RBD. Red lines represent raw data and black lines represent global fit. (E) Comparison of the binding kinetic parameters of 80 as Fab, IgG and MB for binding to WT and Omicron BA.1 RBD. Data shown is average from two independent experiments. (F) Authentic virus neutralization of 80 IgG and MB against wildtype and Omicron BA.2 indicated in grey and red, respectively. The mean values ± SD for two technical replicates is shown in each neutralization plot.

Detailed analysis of the RBD-80 Fab interface revealed that residues S477 and T478 of the RBD form hydrogen bonds with Y^92^ and D^100D^ of the antibody, burying 124 Å^2^ of its surface area and accounting for 15% of the total buried surface area (BSA) of the RBD (Fig. 4B-C, Table S2). These residues are mutated in several VOCs, including Omicron (BA.1, BA.2), which significantly reduces binding affinity of the antibody to the Omicron BA.1 RBD (Fig. 4D-E); however, the increased avidity achieved with the MB format compensates for this weaker binding. Consequently, interaction of 80 MB with the mutated Omicron BA.1 RBD has high apparent binding affinity with no detectable off-rate (Fig. 4D-E), which likely contributes to resilient neutralization potency against Omicron BA.1 (Fig. 3B). The potency of the 80 MB against Omicron BA.2 was additionally confirmed using replication-competent virus: as expected, considerably reduced potency against Omicron BA.2 live virus is observed for the 80 mAb, but high neutralization potency is retained in the MB format (Fig. 4F).

### Broad sarbecovirus neutralization achieved by tri-specific Multabody

Although mono-specific MBs show potent neutralization and can rescue loss in potency compared to their mAb counterparts, mono-specificity still carries the risk of viral escape should sufficient mutations emerge to overcome the benefit conferred by binding avidity. As such, a tri-specific MB targeting three distinct epitopes while retaining avidity has the potential to provide exquisite resilience against evolving variants. mAbs 10-40 and 11-11 have been shown to target conserved epitopes, and consequently confer broad neutralization that expand to other sarbecoviruses^47^. Based on this and the results from the mono-specific MB screening (Fig. 3B), we chose mAb specificities 2-7, 10-40 and 11-11 to design a tri-specific molecule to explore neutralization gains made by combining next-generation mAbs of different epitope specificities on the MB. Similar to mAb 80, previous structural data on mAb 2-7 revealed that RBD mutations found in VOCs form part of its binding interface^50^ (Fig. S8A-B) leading to the neutralization potency loss observed for this IgG against Omicron BA.1. However, 2-7 was rescued in neutralization potency in the MB format (Fig. 5A) similarly to what was observed for mAb 80, demonstrating again the benefit of avidity to overcome viral escape (Fig. S8C-D).

**Figure 5.**
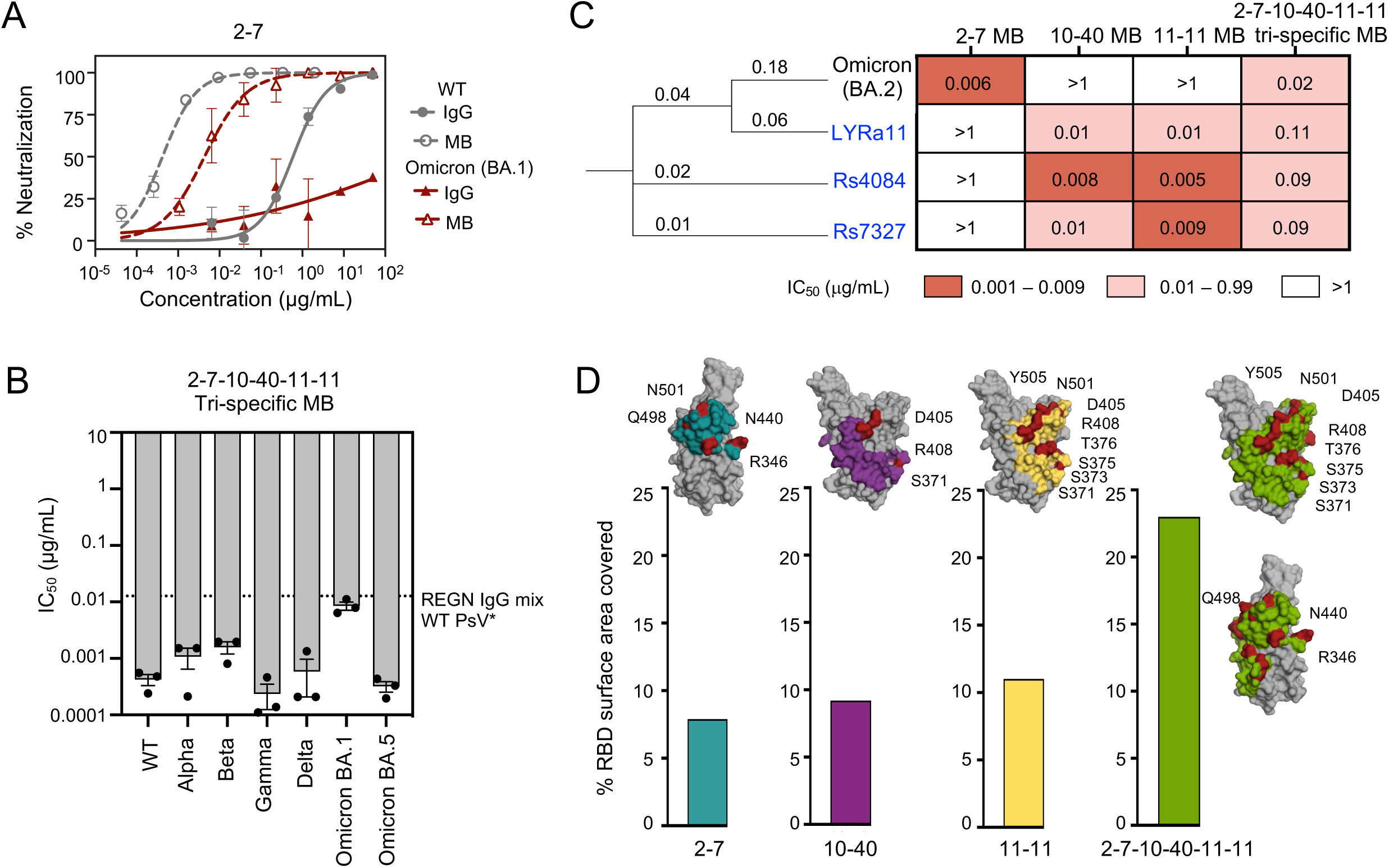
Potent and broad sarbecovirus neutralization by tri-specific Multabody. (A) PsV virus neutralization of 2-7 IgG (solid line) and MB (dashed lines) against WT (gray) and Omicron (BA.1, red). The mean values ± SD for two technical replicates is shown in each neutralization plot. (B) Neutralization potency (grey bars) and breadth of 2-7-10-40-11-11 tri-specific MB against SARS-CoV-2 PsV and six VOCs. *Dashed line indicates IC_50_ of the REGN IgG mix against WT SARS-CoV-2 PsV. (C) Phylogenetic tree with branch lengths representing divergence. Heat map showing neutralization potency of 2-7-10-40-11-11 tri-specific MB and its corresponding mono-specific MBs against Omicron (BA.2) live virus and three SARS-CoV-1 related bat coronaviruses (labeled in blue) PsVs. Individual IC_50_ values are shown. (D) Bar graphs indicating percentage accessible surface area on the RBD (grey) covered by the tri-specific MB (green) and respective components, 2-7 (teal) (PDB ID: 7LSS), 10-40 (purple) (PDB ID: 7SD5), 11-11 (yellow) (EMD: 25167). Mutations found in SARS-CoV-2 VOCs (Alpha, Beta, Gamma, Delta, Omicron (BA.1, BA.2)) that are part of each antibody binding interface are shown in red.

The resulting tri-specific (2-7-10-40-11-11) MB potently neutralized WT and all tested VOCs of SARS-CoV-2 in PsV neutralization assays, with a mean IC_50_ of 0.002 μg/mL (Fig. 5B and Fig. S9A). These experiments were expanded using other viruses dependent on ACE2 as an entry receptor, including live Omicron BA.2 virus and a SARS-CoV-1 related bat sarbecovirus panel. To better assess the benefit of combining multi-specificity, the mono-specific MB components were also tested. Mono-specific 2-7 MB did not show neutralization against the sarbecovirus panel, while 10-40 and 11-11 MBs were not able to block infection of live Omicron BA.2 (Fig. 5C and Fig. S9B-C). In contrast, the tri-specific MB of these specificities combined on a single molecule displayed both potent SARS-CoV-2 neutralization across the VOCs, including live Omicron BA.2, as well as pan-sarbecovirus neutralization of this panel (Fig. 5B-C and Fig. S9A-C).

To gain insight of the molecular coverage of the tri-specific 2-7-10-40-11-11 MB, we analyzed the individual and combined RBD buried surface area of each antibody specificity based on their previously published three-dimensional structures. Binding of mAbs 2-7, 10-40 and 11-11 to RBD covers approximately 8%, 9% and 11% of the RBD accessible BSA, respectively, whereas in the case of the 2-7-10-40-11-11 tri-specific MB, 23% of the RBD accessible BSA is covered with a single molecule (Fig. 5D). Correspondingly, the tri-specific 2-7-10-40-11-11 MB contains 62 contact residues in the RBD compared to 23, 27 and 37 residues in the case of individual mAbs 2-7, 10-40 and 11-11, respectively. Therefore, by targeting three partially overlapping functional epitopes on the RBD, and through the potency gain provided by avidity, the tri-specific 2-7-10-40-11-11 MB provides proof-of-concept for potent, broad, and resilient neutralization across sarbecoviruses by a single molecule.

## Discussion

The rapid emergence of new SARS-CoV-2 VOCs has stymied mAb therapeutics and driven antibody discovery efforts focused on expanding the breadth of viral sequences recognized by a single antibody^21, 22, 51^. The evidence that several FDA-authorized mAb therapies lost efficacy against the Omicron VOC^16, 20^ supports the urgent need for new therapeutic interventions with improved breadth. In addition, strategies to propel the potency of such mAbs have the potential to reduce the therapeutic dose and enable more practical routes of administration, which could reduce manufacturing costs and enable global availability. We have previously described a Multabody platform capable of delivering highly potent and broadly-acting molecules *in vitro*^38, 39^. Indeed, two years after the identification of the tri-specific 298-52-80 MB, derived from three mAbs of modest potency, to our knowledge, no mAb that surpasses the *in vitro* neutralization potency of this molecule against WT SARS-CoV-2 (IC_50_ of 0.0002 µg/mL) has yet been described. Here, we have demonstrated that potent *in vitro* neutralization translates into *in vivo* SARS-CoV-2 protection at low dose and that combination of avidity and multi-specificity can yield molecules with broad neutralization coverage against sarbecoviruses.

The MB platform offers multiple advantages as a next-generation multivalent biologic, including high stability, efficient assembly, ease of production and purification, and plug-and-play genetic fusion of antibodies of choice^38, 39^. Here, we further confirmed the proper assembly of a tri-specific MB by cryoEM. This structural technique has been useful for the characterization of large and complex biological designs such as subunit vaccines, including self-assembling protein nanoparticles presenting the ectodomains of influenza and RSV viral glycoprotein trimers^52^, two-component protein nanoparticles displaying a stabilized HIV-1 Env trimer^53^, or a COVID-19 vaccine candidate nanoparticle utilizing SpyCatcher multimerization of the SARS-CoV-2 spike protein RBD^54^. Even though simultaneous visualization of both nanoparticle scaffold and molecules displayed at their periphery can be challenging due to the flexibility of linkers, recent advances in cryoEM data processing allow for independent analysis of different nanoparticle components^55, 56^. By adopting this strategy, we were able to confirm the proper folding of scFabs and scFcs at the periphery of the MB. Furthermore, we confirm at 2.1 Å resolution the proper assembly of the human light chain apoferritin scaffold of the MB in the context of the engineered split design, on which the assembly of multi-specific antibody components is built. These analyses are important as they further support a central role for structure-guided protein engineering in the rational design of novel biologics and substantiate the MB as a uniform biologic.

Next, we investigated the ability of the MB to confer protection from lethal challenge *in vivo*. The specific role of increased neutralization potency in mediating *in vivo* protection was assessed with a tri-specific MB* molecule expressing a mutant IgG4 Fc that is defective for Fcγ receptor binding. In attempts to retain IgG-like bioavailability as previously described^38^, we maintained the ability of the MB Fc to interact with FcRn, a receptor associated with antibody recycling and half-life extension^57, 58^. *In vivo*, the dramatic increase in neutralization potency of the tri-specific MB relative to the IgG4* cocktail resulted in significantly better protection against lethal SARS-CoV-2 challenge, and facilitated a reduction in the dose required for protection. Previous studies have shown that some mAbs targeting SARS-CoV-2 require Fc-mediated effector functions for optimal efficacy^59–63^. Our data illustrates that gains in neutralization potency are sufficient to confer protection from lethal challenge, even in the absence of effector function, and supports the use of avidity-based increases in potency to facilitate dose-sparing of antibody-based therapeutics. Further studies will investigate whether MB potency can be further enhanced through the addition of effector functions, by incorporating WT Fc or engineering the Fc to introduce specific functions. We have previously shown that the MB format is capable of triggering ADCP *in vitro,* indicating that the format in itself does not preclude incorporating effector function into the molecule^39^.

Single-specificity MBs showed an elevated degree of resilience against viral sequence variability through improvements in their apparent binding affinity compared to IgGs, allowing these molecules to retain neutralization capabilities even when mAbs lose potency. In the case of both mAbs 80 and 2-7, mutations within the RBD epitope found in the VOCs cause a loss in potency by the mAbs. In contrast, when both specificities are displayed as MBs, mutations present in the VOCs minimally alter the high binding affinity and potent neutralization profiles of these molecules. The ability of the MB to better tolerate sequence variability presumably stems from the reduced off rate that drives increased avidity, which might be favoured by the spike density on the virion surface^64–66^. The potential for avid antibody technologies to endure viral sequence variability can provide benefits to antibody discovery timelines by boosting the longevity of early identified mAbs with the ability to neutralize emerging VOCs.

The identification of potent bnAbs can take years or even decades of antibody discovery and engineering efforts, as exemplified in the cases of HIV-1 and Influenza. Here, we demonstrate that a tri-specific MB incorporating the specificities 2-7, 10-40, and 11-11 has potent *in vitro* neutralization across all SARS-CoV-2 VOCs tested and expand its neutralization breadth to other sarbecoviruses at neutralization potencies within the range of FDA-authorized therapeutics against SARS-CoV-2^49^. The continuous monitoring and screening of emerging variants will be required to confirm persistence in neutralization; however, the larger footprint of the RBD covered by a tri-specific MB compared to conventional mAbs, provides a unique advantage for the MBs in remaining resilient against future VOCs compared to mAbs alone. Furthermore, the ability to combine multiple specificities into a single molecule might offer the additional potential benefit of ensuring the bioavailability of all components throughout the course of therapy, which has been a limiting feature of mAb cocktail combinations^70, 71^. Further studies of the pharmacokinetics of the MBs compared to corresponding cocktails of IgGs in higher-order species will evaluate this potential benefit. More broadly, ongoing efforts will explore the potential of combining the most potent bnAbs that continue to emerge from discovery efforts on an avid antibody platform to evaluate the full potential of this approach against betacoronaviruses. The MB platform presents a promising avenue to explore also in the context of other viral pathogens of high genetic diversity such as HIV in attempts to further propel the breadth and potency of antibody therapeutics against indications of global health importance.

## Materials and Methods

### Cell lines and viruses

HEK 293T-ACE2 cells (BEI NR2511), HEK 293T (ATCC) and Vero E6 (Green Monkey Kidney cell line, ATCC) cells were grown in DMEM (Thermo Fisher Scientific, Waltham, MA). For HEK 293T and 293T-ACE2 cells, DMEM was supplemented with 10% heat inactivated FBS, 5% 1M HEPES and 1% gentamicin (all Thermo Fisher Scientific, Waltham, MA). For Vero E6 cells, DMEM was supplemented with 10% FBS (Hyclone, Logan, UT) and 1% penicillin-streptomycin (Invitrogen, Thermo Fisher Scientific, Waltham, MA). HEK 293F and HEK 293S cells (Thermo Fisher Scientific, Waltham, MA) were cultured in Freestyle 293 Expression Medium (Thermo Fisher Scientific, Waltham, MA) at 125 rpm oscillation, 37 °C, 8% CO_2_. SARS-CoV-2/SB2-P4-PB Clone 1^69^ titers were determined by 50% tissue culture infectious dose (TCID50/mL) using cell supernatants as previously described^70, 71^.

### Protein expression and purification

Genes encoding human apoferritin fusions, Fabs, Fcs and IgGs were synthesized and cloned by GeneArt (Thermo Fisher Scientific, Waltham, MA) in the pcDNA3.4 expression vector and transiently expressed in HEK 293F cells. IgG1 and IgG4* versions of Fc were used, where IgG4* indicates the inclusion of the following mutations in the IgG4 Fc to ablate Fcγ receptor binding: S228P, F234A, L235A, G237A, P238S^43^. Cells were cultured at a density of 0.8 x 10^6^ cells/mL and transfected with 50 µg of DNA per 200 mL of cells using FectoPRO (Polyplus Transfections, Strasbourg, France) as previously described^38^. Following 6-7 days of incubation with oscillation (Multitron Pro shaker, Infors HT, 125 rpm oscillation, 37 °C, 8% CO_2_, 70% humidity), cells were harvested by centrifugation at 5000 rpm for 20 min and supernatants filtered through a 0.22 µm Steritop filter (EMD Millipore, Burlington, MA). Fabs and IgGs were expressed by transiently co-transfecting 90 µg of heavy and light chain at a 2:1 ratio, and purified using KappaSelect affinity and HiTrap Protein A HP columns, respectively (both GE Healthcare, Chicago, IL), eluted with 100 mM glycine pH 2.2, and neutralized with 1 M Tris-HCl, pH 9.0. IgG fractions were further purified by size exclusion chromatography (Superdex 200 Increase, GE Healthcare, Chicago, IL), and Fab fractions were further purified by cation exchange chromatography (MonoS, GE Healthcare, Chicago, IL). His-tagged wildtype RBD (BEI NR52309), human and mouse Fc receptors (hFcγRI, hFcγRIIa, hFcγRIIb, hFcRn, mFcγRI, mFcγRIIb, mFcγRIV, mFcRn) were purified using a HisTrap Ni NTA column followed by size exclusion chromatography (Superdex 200 Increase column; both GE Healthcare, Chicago, IL) using 20 mM phosphate pH 8.0, 150 mM NaCl buffer.

### Expression and purification of Multabodies

MBs were designed, expressed and purified as previously described^38^. Briefly, genes encoding single chain (sc) Fab and scFc linked to human apoferritin light chain monomers were synthesized and cloned by Geneart (Thermo Fisher Scientific, Waltham, MA) into the pcDNA3.4 expression vector. MBs were expressed by transient transfection of 66 µg of plasmid (scFab-apoferritin: scFc-N-Ferr: scFab-C-Ferr at a 2:1:1 ratio for mono-specific MBs; or a 4:2:1:1 ratio of scFab1-apoferritin: scFc-N-Ferr: scFab2-C-Ferr: scFab3-C-Ferr for tri-specific MBs) in to HEK 293F cells using FectoPRO (Polyplus Transfections, Strasbourg, France). MBs were purified by affinity chromatography using a HiTrap Protein A HP column (GE Healthcare, Chicago, IL) and eluted with 20 mM Tris pH 8.0, 3 M MgCl_2_ and 10% glycerol. Fractions were concentrated and further purified by gel filtration (Superose 6 10/300 GL column, GE Healthcare, Chicago, IL). For the tri-specific MB (298-52-80), an IgG1 or IgG4* scFc was used, where MB* indicates the use of an IgG4 with the Fcγ receptor binding mutations specified above^43^. For *in vivo* studies, all IgGs and MBs were quality controlled to ensure endotoxin levels below 3.5 EU/mL at a 1 mg/mL concentration of protein^72^.

### CryoEM data collection and image processing

The tri-specific MB (298-52-80) sample was concentrated to 2.0 mg/mL and 3.0 µl of the sample was deposited on homemade holey gold grids^73^, which were glow-discharged in air for 15 s before use. Sample was blotted for 3.0 s, and subsequently plunge-frozen in liquid ethane using a Leica EM GP2 Automatic Plunge Freezer (maintained at 4 °C and 100% humidity). Data collection was performed on a Thermo Fisher Scientific Titan Krios G3 operated at 300 kV with a Falcon 4i camera automated with the EPU software. A nominal magnification of 75,000× and defocus range between 0.5 and 2.0 μm were used for data collection. Exposures were collected for 8.3 s as movies of 30 frames with a camera exposure rate of ∼6.3 e^−^ per pixel per second, and total exposure of 49.6 electrons/Å^2^. A total of 4,385 raw movies were obtained.

Image processing was carried out in cryoSPARC v3^74^. Initial specimen movement correction, exposure weighting, and CTF parameters estimation were done using patch-based algorithms. Micrographs were sorted based on CTF fit resolution, and only micrographs with a fit better than 5.0 Å were accepted for further processing. Manual picking was performed to create templates for template-based picking, which resulted in selection of 955,995 particle images. Particle images were sorted via several rounds of 2D classification, which resulted in selection of 358,036 particle images. A preliminary 3D model was obtained *ab-initio* with no symmetry applied. To further select the highest-quality particle images, 151,443 particle images with CTF fit resolution better than 3.0 Å were re-extracted from micrographs and subjected to non-uniform refinement^75^ with no symmetry applied, which resulted in a 2.4 Å resolution map of the tri-specific MB. 65,478 particle images with CTF fit better than 2.7 Å, were extracted from micrographs and subjected to non-uniform refinement with octahedral symmetry applied, which resulted in a 2.1 Å resolution map. Non-uniform refinements were performed with defocus refinement and optimization of per-group CTF parameters. The pixel size was calibrated at 1.04 Å per pixel by fitting a structure of human apoferritin light chain (PDB ID: 2FFX)^76^.

To obtain 3D reconstructions of Fab and Fc molecules on the surface of the tri-specific MB, manual picking was performed, and templates were created for template-based picking, resulting in the selection of 6,692,141 particle images. Particle images were sorted via several rounds of 2D classification, which resulted in the selection of 668,214 particle images. Preliminary 3D maps were obtained *ab-initio* with no symmetry applied. Further cleaning of the dataset was performed via several rounds of heterogenous refinement and resulted in 73,163 Fab particle images and 13,328 Fc particle images. Final cryoEM maps at 6.7 Å resolution for Fab and 7.1 Å resolution for Fc were obtained using the local refinement job with a custom soft mask.

To assess the quality of obtained maps, models of human apoferritin light chain (PDB ID: 6WX6)^77^, human IgG1 Fc (PDB ID: 6CJX)^78^ and Fab 298 (PDB ID: 7K9Z)^38^ were manually docked in cryoEM maps using UCSF Chimera^79^. Figures were prepared with Pymol^80^, UCSF Chimera^79^ and UCSF ChimeraX^81^.

### Pseudovirus production

Pseudoviruses (PsV) were produced using a lentiviral vector backbone, as previously described^82^. Briefly, SARS-CoV-2 PsV were generated by transient co-transfection of 293T cells with a lentiviral backbone encoding luciferase reporter gene (BEI NR52516), plasmids encoding Gag-Pol (BEI NR52517), Tat (BEI NR52518), Rev (BEI NR52519) and a plasmid expressing SARS-CoV-2 Spike (BEI NR52310) with BioT transfection reagent (Bioland Scientific, Paramount, CA) according to the manufacturer’s directions. Cells were incubated at 37 °C for 24 h, followed by addition of 5 mM of sodium butyrate. Cells were further incubated for an additional 24-30 h at 30 °C. Omicron (BA.1) variant PsV was generated using the Omicron (BA.1) Spike plasmid kindly provided by D.R. Burton (Scripps Research). Genes encoding the Omicron (BA.5) Spike was synthesized and cloned by GeneArt (Thermo Fisher Scientific, Waltham, MA) in the pcDNA3.4 expression vector. Alpha, Beta, Gamma, Delta, and Omicron SARS-CoV-2 PsV variants were generated by substituting the WT Spike plasmid. PsV were harvested, filtered through 0.45 µm sterile filters, and concentrated using a 100 K Amicon filter (Merck Millipore Amicon – Ultra 2.0 Centrifugal Filter Units, Millipore Sigma, Burlington, MA).

### PsV neutralization assay

Single-cycle neutralization using 293T-ACE2 cells (BEI NR52511) was used to assess neutralization, as previously described^38^. Briefly, 96-well plates were coated with poly-L-Lysine (Sigma Aldrich, St. Louis, MO) and cells were seeded at a density of 10,000 cells/well in 100 μL. The following day, IgGs or MBs were serially diluted in duplicate and incubated with PsV for 1 h at 37 °C. Cell culture medium was then replaced with the PsV-IgG or PsV-MB mixture supplemented with 10 µg/mL polybrene (Sigma-Aldrich, St. Louis, MO). Cells were incubated for 48 h at 37 °C, before addition of 50 μL of Britelite plus reagent (PerkinElmer, Waltham, MA) for 2 min. Supernatants were transferred to 96-well white plates (Sigma Aldrich, St. Louis, MO) and luminescence in relative light units (RLUs) was measured using a Synergy Neo2 Multi Mode Assay Microplate Reader (Biotek Instruments, Winooski, VT). Three biological replicates were performed for each antibody or MB. IC_50_ values were calculated using nonlinear regression in GraphPad Prism 9.3.1.

### Pseudovirus production of sarbecoviruses

Spike gene sarbecovirus S genes were codon-optimized for mammalian expression, synthesized by Twist Biosciences, and cloned into the same expression vectors as above by Gibson Assembly (New England Biolabs). Sarbecovirus sequences were retrieved from GenBank for Rs4084, Rs7327, and LYRa11 as shown in Table S3. Recombinant VSV pseudoviruses in which the native glycoprotein was replaced with sarbecovirus S proteins were generated as previously described^47, 82^. Briefly, human embryonic kidney (HEK) 293T cells (ATCC), at a confluency of 80% were transfected with a S protein expression vector using PEI (1 mg/ml) and cultured overnight at 37 °C under 5% CO_2_. Twenty-four hours later, cells were infected with VSV-G– pseudotyped ΔG-luciferase (G*ΔG-luciferase, Kerafast) at a multiplicity of infection (MOI) of 3 for 2 h. Following infection, cells were washed three times with 1× PBS, changed to fresh medium, and cultured at 37 °C for another 24 h before supernatants were harvested and clarified by centrifugation at 300*g* for 10 min.

### Sarbecovirus neutralization assay

Pseudoviruses were titrated to standardize the infectivity levels for target cells before setting up neutralization assays. Neutralization assays were performed as described earlier by incubating pseudoviruses with 5-fold serial dilutions of Multabody versus their corresponding IgGs in triplicate in a 96-well plate for 1 h at 37 °C^47^. Briefly, 293T-hACE2 cells were seeded at a density of 1 × 10^5^ cells per well. Luciferase activity was measured using the Luciferase Assay System (Promega), according to the manufacturer’s instructions, 24 h after cells were added to the pseudovirus and serum. The neutralization curves and IC_50_ values were generated by fitting a nonlinear five-parameter dose-response curve in GraphPad Prism 9.3.

### Virus propagation and titration

The SARS-CoV-2 Omicron isolate hCoV-19/USA/MD-HP24556/2022 (BA.2) was obtained from BEI Resources (NIAID, NIH). The virus was propagated using Vero E6 cells. Virus infectious titer was determined by an end-point dilution and cytopathic effect (CPE) assay on Vero E6 cells as described previously^24, 47, 83^.

An end-point-dilution microplate neutralization assay was performed to measure the neutralization activity of IgG and Multabodies. Triplicates of each dilution were incubated with SARS-CoV-2 at an MOI of 0.1 in EMEM with 7.5% inactivated fetal calf serum (FCS) for 1 h at 37 °C. Post incubation, the virus-antibody mixture was transferred onto a monolayer of Vero E6 cells grown overnight. The cells were incubated with the mixture for ∼70 h. CPE was visually scored for each well in a blinded fashion by two independent observers. The results were then converted into percentage neutralization at a given sample dilution, and the averages ± SEM were plotted using a five-parameter dose-response curve to obtain the ID50 of each sample using GraphPad Prism v.9.3.

### Biolayer interferometry

Binding kinetics were measured using an Octet RED96 Biolayer Interferometer (Sartorius ForteBio, Freemont, CA)^30^. His-tagged Omicron RBD (SinoBiological, China), wildtype RBD, human or mouse Fc gamma receptors (hFcγRI, hFcγRIIa, hFcγRIIb, mFcγRI, mFcγRIIb, mFcγRIV) was loaded on to Ni-NTA biosensors (Sartorius ForteBio, Freemont, CA) to reach a 0.8 nm signal response. Association rates were measured by transferring the loaded sensors to wells containing either Fab (titrated from 150 nM to 4.7 nM), IgG (titrated from 150 nM to 4.7 nM) or MB (titrated from 20 nM to 0.6 nM) for 180 s. Dissociation rates were measured by dipping the sensors into buffer-containing wells for 180 s. All steps were performed in buffer containing PBS pH 7.4, 0.01% BSA and 0.0002% Tween-20 at 25 °C. To estimate the potential for endosomal recycling, binding of IgG or MB to mouse and human neonatal Fc receptor (FcRn) was assessed. IgGs and MBs were titrated at the concentrations listed above. All binding and incubation steps except dissociation were performed in buffer containing PBS pH 5.6 and 0.0002% Tween. Dissociation step was measured using PBS pH 7.4 and 0.0002% Tween-20. Analysis was performed using the Octet software, with a 1:1 fit model.

### Preparation of SARS-CoV-2 Spike microspheres

SARS-CoV-2 Spike protein (Sino Biological, Beijing, China) was biotinylated using the EZ-link Sulfo-NHS biotinylation kit (Thermo Scientific, 2143) according to the protocol provided. Red fluorescent Neutravidin microspheres (Invitrogen, F8775) were washed twice with PBS + 0.1% BSA before incubating 5 µL with 10 µg of biotinylated protein and bringing the total volume to 200 µL with PBS/0.1% BSA. Beads were incubated overnight at 4 °C and washed twice before use to remove unbound protein. Beads were resuspended in 200 µL per 5 µL bead volume.

### Antibody-dependent cell-mediated phagocytosis assay

Immune complexes were formed by incubating SARS-CoV-2 Spike-coated fluorescent beads with diluted MB or IgG preparations for 2 h at 37 °C + 5% CO_2_ (10 µL beads and 10 µL of 1 mg/mL antibody sample). THP-1 cells (ATCC, TIB-202) were maintained at fewer than 5 x 10^5^ cells/mL and 5 x 10^4^ cells/well in 200 µL were added to the immune complexes for 1 h at 37 °C + 5% CO_2._ Cells were washed and stained with Live Dead Fixable Violet stain (Invitrogen, L34995) according to the provided protocol before being washed and fixed with 1% PFA for 20 min at room temperature. Fixed cells were washed with FACS buffer (PBS + 10% FBS, 0.5 mM EDTA) and collected on an LSRII Flow Cytometer (BD Biosciences). Data was analyzed in FlowJo (BD Biosciences, Ashland, OR), and phagocytosis was quantified as the percentage of live THP-1 cells that had phagocytosed red fluorescent SARS-CoV-2 Spike beads.

### Mice and ethics statement

Age-matched female human ACE2-expressing and human FcRn-expressing mice (JAX strain #034902, B6.Cg-Tg(FCGRT)32Dcr Tg(K18-ACE2)2Prlmn Fcgrttm1Dc) were purchased from the Jackson Laboratory (JAX, Bar Harbor, ME) and housed in individually-ventilated caging under specific pathogen-free conditions. All procedures were approved by the Local Animal Care Committee at the University of Toronto, AUP#20012628. Studies with replication competent SARS-CoV-2 were performed according to the University of Toronto’s Containment Level 3 guidelines under Biosafety Permit EXT-J06-3.

### SARS-CoV-2 challenge

A pre-challenge oropharyngeal swab (Good Care nasopharyngeal swabs, Goodwood Medical Care, Dalian, China) sample was collected and female B6.Cg-Tg(FCGRT)32Dcr Tg(K18-ACE2)2Prlmn Fcgrttm1Dc mice were treated intraperitoneally (i.p) with 3 µg (0.15 mg/kg), 30 µg (1.5 mg/kg) or 90 µg (4.5 mg/kg) of cocktail IgG4*; tri-specific MB* (298-52-80) or a negative control (PGDM1400 IgG) as specified before transfer to the Combined Containment Level 3 *in vivo* facility at the University of Toronto for SARS-CoV-2 challenge. Mice were anesthetized with inhaled isoflurane and inoculated intranasally with a lethal dose of 1 x 10^5^ pfu/mouse of SARS-CoV-2 (SB2-P4-PB Clone 1)^69^. Mice received a second dose of tri-specific MB* or IgG4* cocktail one day following challenge; negative control recipient animals also received a second 30 µg dose of PGDM in dose investigation studies. Oropharyngeal swabs were collected two days post-challenge and subject to qPCR quantification to validate infection. Mice were weighed and monitored daily and scored according to weight loss and disease progression. Clinical disease was scored as follows: 0) no signs of disease; 1) 5 – 10% weight loss with no additional symptoms; 2) slightly reduced movement and 10 – 20% weight loss; 3) reduced movement or limited unprovoked movement with < 20% weight loss; 4) lack of provoked movement, rapid breathing, hunching and reduced grooming; or body weight loss of 20% or more; 5) Death. Mice were sacrificed at a score of 4 and given a score of 5 the following day.

### Lung tissue viral outgrowth

At the time of sacrifice or 12 days following challenge for surviving animals, lung tissue was collected and snap frozen. Lung viral titer at the time of sacrifice was determined by homogenizing lung tissue in 0.5 mL of DMEM (Invitrogen, Carlsbad, CA) without additives, and calculating TCID50 on Vero E6 cells by performing ten-fold serial dilutions and monitoring cytopathic effect (CPE)^69–71^.

### Enzyme-linked immunosorbent assay

Serum collected at D2 following molecule administration was diluted 1:100 and evaluated for MB or antibody concentration by ELISA. Briefly, Nickel-coated 96-well plates (Pierce, Thermo Fisher Scientific, Waltham, MA) were coated overnight with 50 µL per well of His-tagged SARS-CoV-2 RBD protein or, for PGDM1400, BG505 D368R SOSIP.664 trimer^39^. Plates were blocked with TBS + 5% BSA for 1 h at room temperature and incubated with diluted serum or a standard curve of each of the administered molecules. MB and antibody were detected using an anti-human Fab secondary antibody (Abcam, 87422) and developed using BD TMB substrate reagent set (BD Biosciences, Franklin Lakes, NJ). Data was collected using a Synergy Neo2 Multi-Mode Assay Microplate Reader (Biotek Instruments, Winooski, VT).

### qPCR for oropharyngeal swab titers

Viral RNA from swab samples were extracted using a QIAamp Viral RNA Mini Kit (Qiagen, Hilden, Germany). Primers targeting *env* (Forward Primer: ACAGGTACGTTAATAGTTAATAGCGT, Reverse Primer: ATATTGCAGCAGTACGCACACA) as described by Corman *et al*^84^ were used alongside the Luna Universal One-Step RT-qPCR kit (New England Biolabs, Ipswitch, MA) and CFX384 Touch Real-Time System (Bio-Rad) for RT-qPCR. Nuclease-free water was used as a no template control. Reverse transcription was conducted by incubation for 10 min at 55 °C and PCR cycle conditions consisted of 1 min at 95 °C for initial denaturation, followed by 45 cycles of denaturation (95 °C for 10 s) and extension (58°C for 30 s). The melting curve was generated by running 5 s cycles between 65 °C to 95 °C at intervals of 0.5 °C. A reference copy number plasmid for *env*-gene was used to generate a standard curve. Data was analyzed using CFX Maestro Software (Bio-Rad).

### Crystallization and structure determination

A binary complex of purified 80 Fab-RBD was obtained by mixing Fab:RBD in a 2:1 molar ratio. After 30 min incubation at 4 °C, the complex was purified by size exclusion chromatography (Superdex 200 Increase size exclusion column, GE Healthcare, Chicago, IL) in 20 mM Tris pH 8.0, 150 mM NaCl buffer. The fractions of interest were then concentrated to 10 mg/mL and crystallization trials were set up using the sitting drop vapor diffusion method with JCSG Top 96 screen in a 1:1 protein: reservoir ratio. Crystals appeared on day 70 in a condition containing 0.2 M di-ammonium tartrate and 20% (w/v) PEG 3350. Crystals were cryoprotected in 10% (v/v) ethylene glycol and flash-frozen in liquid nitrogen. X-ray diffraction data was collected at the Argonne National Laboratory Advanced Photon Source on the 23-ID-D beamline. The data set was processed using XDS^85^ and XPREP^86^. Phases were determined using Phaser^87^ with the 80 Fab predicted by ABodyBuilder^88^ and the SARS-CoV-2 RBD (PDB ID: 7LM8)^89^ as search models. Iterative refinement was performed using Phenix Refine^90^ and manual building was done in Coot^91^. All software were accessed through SBGrid^92^. Modeling of 2-7 Fab binding to Omicron (BA.1, BA.2) was performed in Pymol^80^ using the RBD structure published by Cerutti *et al* (PDB ID: 7LSS)^50^. Footprint analysis was done using the 2-7 Spike complex (PDB ID: 7LSS)^50^, 10-40 RBD complex and 11-11 Spike complex structures published by Liu et al (PDB ID:7SD5 and EMD-25167). Interface residues were identified with PISA^93^ using default parameters and Pymol^80^ was used to generate figures.

### Statistical analysis

Statistical analyses were performed using Graphpad Prism version 9.3.1 software (Graphpad Software Inc., San Diego, CA). A P value of 0.05 was considered statistically significant, unless adjusted for multiple testing using the Bonferroni correction. Survival curves were compared using the Gehan-Breslow-Wilcoxon test; ADCP effector function data was compared by ANOVA; and viral outgrowth data (not normally distributed) was compared using the Kruskal-Wallis test. Data are shown as means ± SEM unless otherwise indicated.

### Data availability

The electron microscopy maps have been deposited in the Electron Microscopy Data Bank (EMDB) with accession codes EMD-28067 (tri-specific MB, refinement with no symmetry) and EMD-28068 (tri-specific MB, octahedral symmetry). The crystal structure of 80 Fab-RBD has been deposited in the Protein Data Bank (PDB ID: 8DNN).

## Acknowledgements

We thank A. Banerjee and K. Mossman for their contributions to the initial isolation of SARS-CoV-2 (SARS-CoV.2/SB-2-P4-PB Clone1); F. Krammer for providing the WT S plasmid; D.R. Burton for providing the Omicron BA.1 S plasmid; J.D. Bloom and A.C. Gingras for 293T-ACE2 cells and reagents to make SARS-CoV-2 PsV; and S. Benlekbir for advice regarding cryoEM data collection and specimen preparation. The following reagent was produced under HHSN272201400008C and obtained through BEI Resources, NIAID, NIH: Vector pCAGGS Containing the SARS-Related Coronavirus 2, Wuhan-Hu-1 Spike Glycoprotein Gene, NR-52310. Omicron BA.2 (NR-56512) used for this study was also sourced from BEI Resources, NIAID, NIH.

## Declaration of interests

### Author contributions

C.B.A., K.M., A.J., and J.-P. J conceived the research and designed the experiments; C.B.A., K.M., I.K., H.C., K.P., M.S.N., M.W., Y.H., B.P., J.L., and A.S performed experimental work. N.C.-H, R.K. and S.M. provided critical reagents. N.C.-H., R.K., S.M. and E.R., provided critical expertise. C.B.A., K.M., I.K., M.S.N, M.W., Y.H, J.L.R., B.T., D.D.H., A.J., and J.-P.J. analyzed the data. C.B.A., K.M., I.K., E.R., A.J., and J.-P.J. wrote the manuscript with input from all authors.

---

**Supplementary Fig 1.**
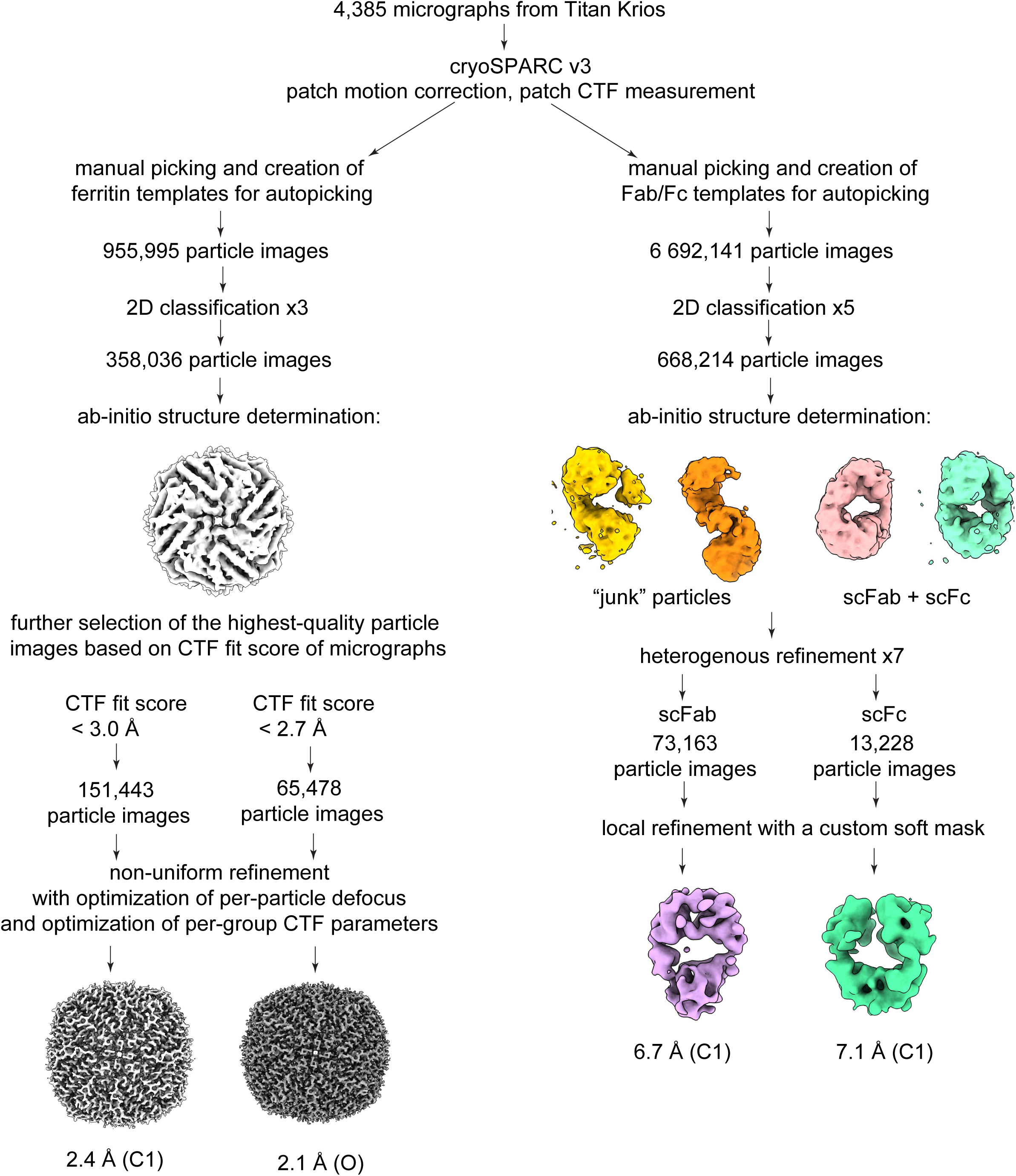
CryoEM data processing workflow in cryoSPARC v3.

**Supplementary Fig 2.**
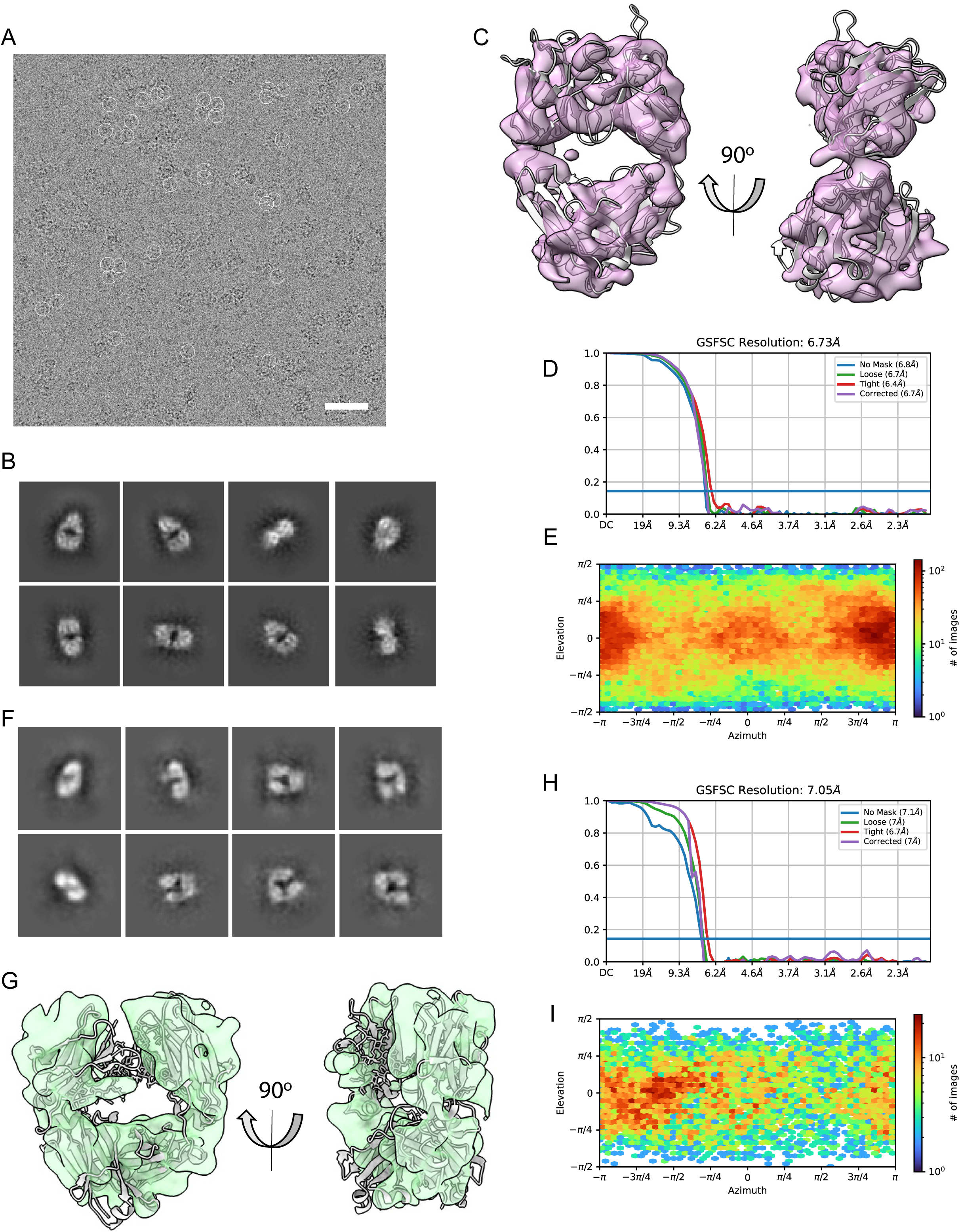
CryoEM analysis of Fab and Fc from the periphery of the tri-specific MB (298-52-80). (A) An example of cryoEM micrograph with Fab and Fc molecules highlighted with white circles. Scale bar is 50 nm. (B) Selected 2D class averages of Fab. (C) Atomic model of Fab 298 (PDB ID: 7K9Z) fit into cryoEM map of Fab. (D) GSFSC curve of the final 3D non-uniform refinement of Fab in cryoSPARC v3. (E) Viewing direction distribution of the Fab data. (F) Selected 2D class averages of Fc. (G) Atomic model of human IgG1 Fc (PDB ID: 6CJX) fit into cryoEM map of Fc. (H) GSFSC curve of the final 3D non-uniform refinement of Fc in cryoSPARC v3. (I) Viewing direction distribution of the Fc data.

**Supplementary Fig 3.**
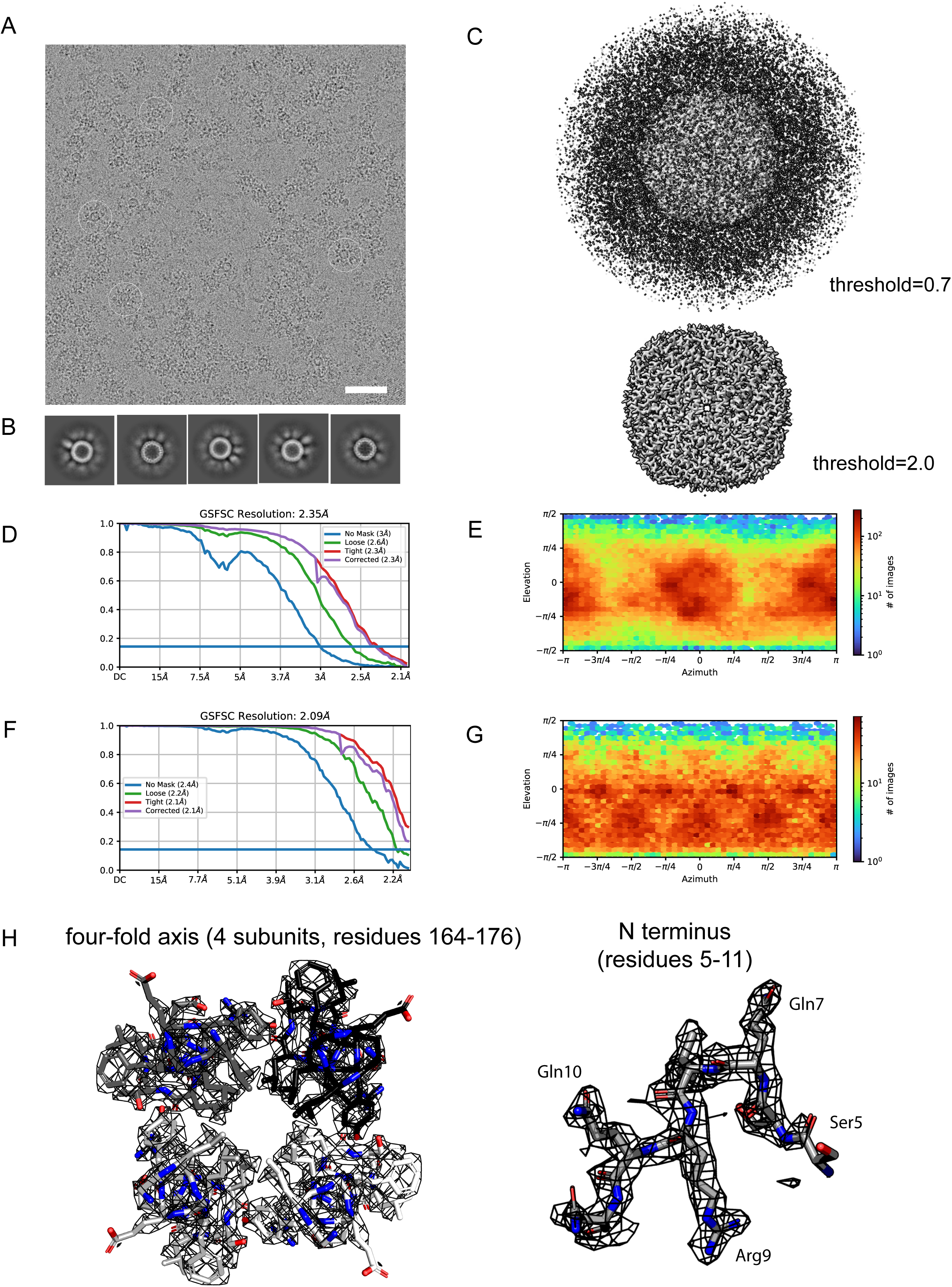
CryoEM analysis of tri-specific MB (298-52-80). (A) Representative cryoEM micrograph of the tri-specific MB particles (highlighted with white circles). Scale bar is 50 nm. (B) Selected 2D class averages of the tri-specific MB particles. (C) Comparison of tri-specific MB (298-52-80) cryoEM reconstruction at two threshold levels (top – 0.7 and bottom – 2.0) reveals weak and fragmented densities for the antibody fragments fused to the apoferritin scaffold. (D) Gold standard Fourier shell correlation (GSFSC) curve of the final 3D non-uniform refinement of the tri-specific MB in cryoSPARC v3 without symmetry imposed. (E) Viewing direction distribution of tri-specific MB dataset refined without symmetry imposed. (F) GSFSC curve of the final 3D non-uniform refinement of the tri-specific MB in cryoSPARC v3 with octahedral symmetry imposed. (G) Viewing direction distribution of the tri-specific MB dataset refined with octahedral symmetry imposed. (H) Details of atomic model of human apoferritin light chain (PDB ID: 6WX6) fit into the tri-specific MB (298-52-80) map refined with octahedral symmetry imposed. CryoEM density of the tri-specific MB is shown as black mesh with the models of adjacent apoferritin protomers shown as white, grey or black sticks.

**Supplementary Fig 4.**
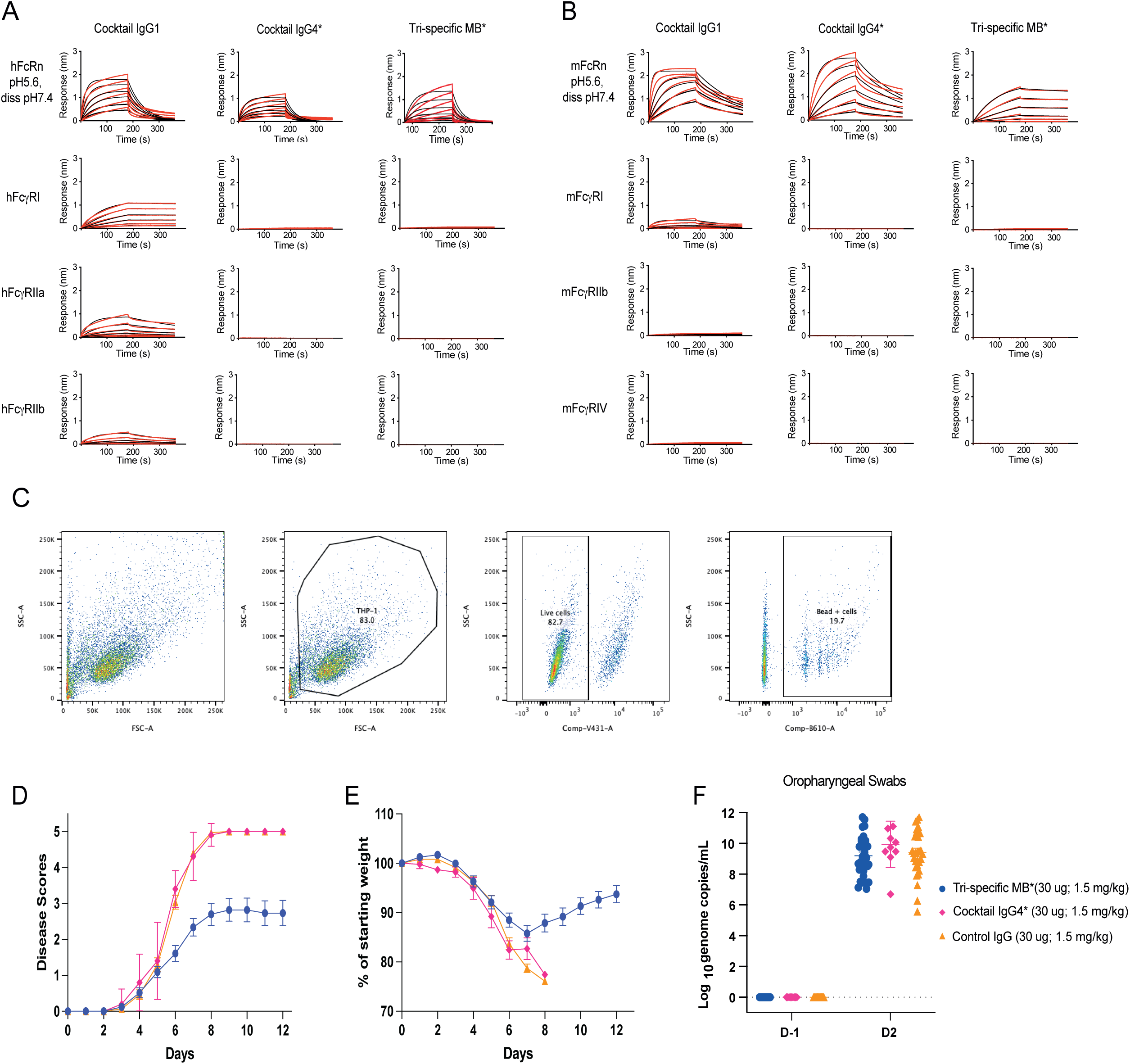
FcγR binding, ADCP and validation of *in vivo* infections. Sensograms of tri-specific MB* and cocktail IgG1 and IgG4* binding to human (A) and mouse (B) Fc receptors. * indicates the use of an IgG4* Fc bearing the specified mutations (PAAAS) to ablate Fcγ receptor binding. Red lines represent raw data and black lines represent global fit. Representative data from 2-3 independent experiments is shown. (C) Gating strategy followed in the ADCP assay. THP-1 cells were gated by size and live cells, and cells positive for internalization of SARS-CoV-2 Spike-coated fluorescent beads were quantified as a percentage of live THP-1 cells. (D) Signs of disease and (E) weight loss were monitored for 12 days following challenge. (D-E) n = 33 for tri-specific MB*, n = 30 for control IgG, n = 10 for cocktail IgG4*, from 2-6 independent experiments. (F) Infection following intranasal SARS-CoV-2 administration was confirmed by collecting oropharyngeal swab samples at D-1 (before challenge) and D2 following challenge. SARS-CoV-2 genome copy number was quantified by qPCR.

**Supplementary Figure 5.**
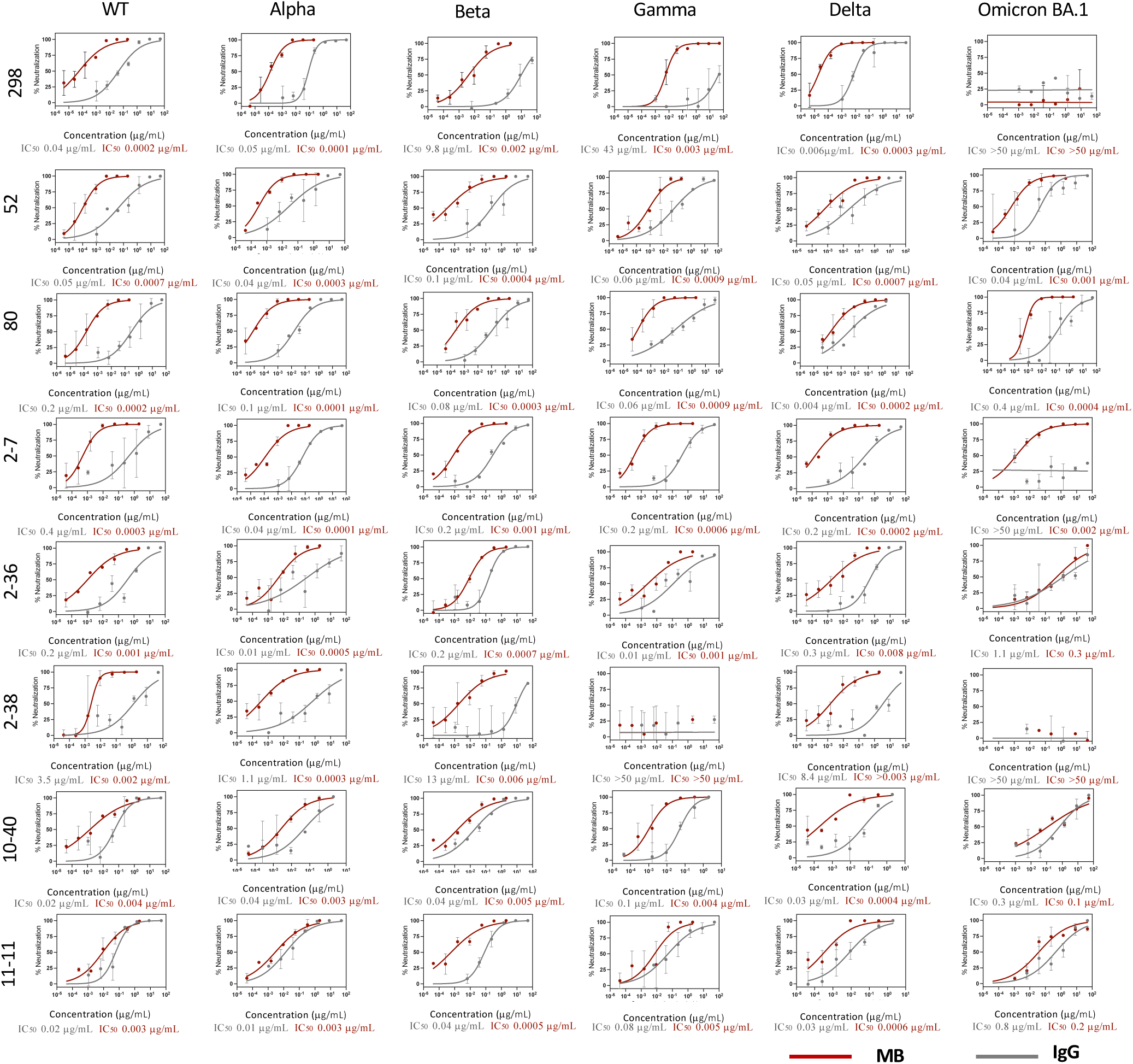
Neutralization of mAbs and corresponding mono-specific Multabodies. Representative neutralization plots of RBD directed IgGs (grey) and corresponding MBs (red) against SARS-CoV-2 WT, Alpha, Beta, Gamma, Delta and Omicron (BA.1) PsVs. The mean values ± SD for two technical replicates is shown in each plot. Mean IC_50_ values from three biological replicates are shown.

**Supplementary Figure 6.**
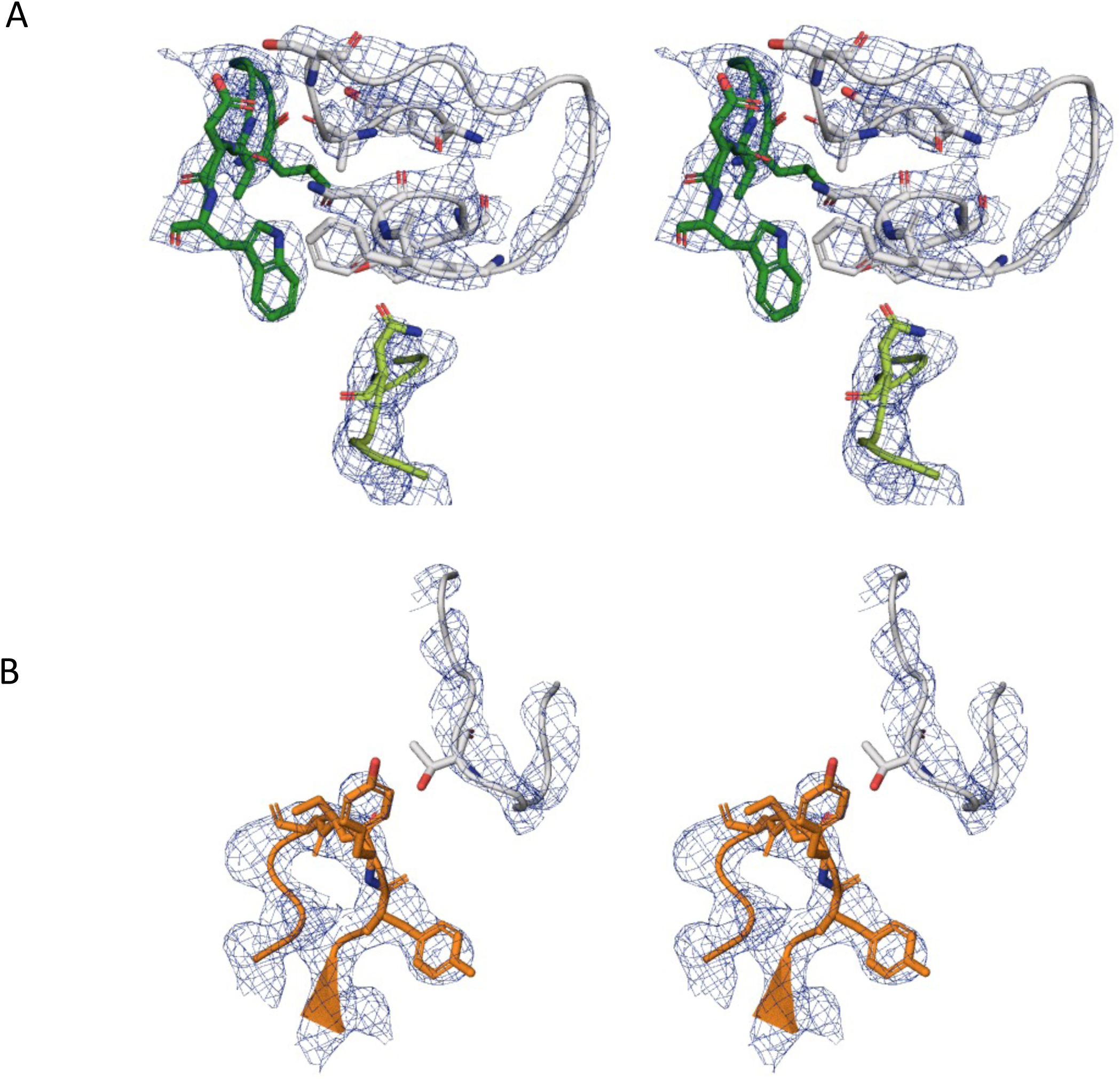
Stereo-image of composite omit map electron density of 80 Fab– RBD interaction sites. (A) Heavy chain complementarity determining regions (HCDRs) 2 and 3 (light and dark green, respectively) and RBD (grey). (B) Kappa light chain CDR (KCDR) 3 (orange) and RBD (grey).

**Supplementary Figure 7.**
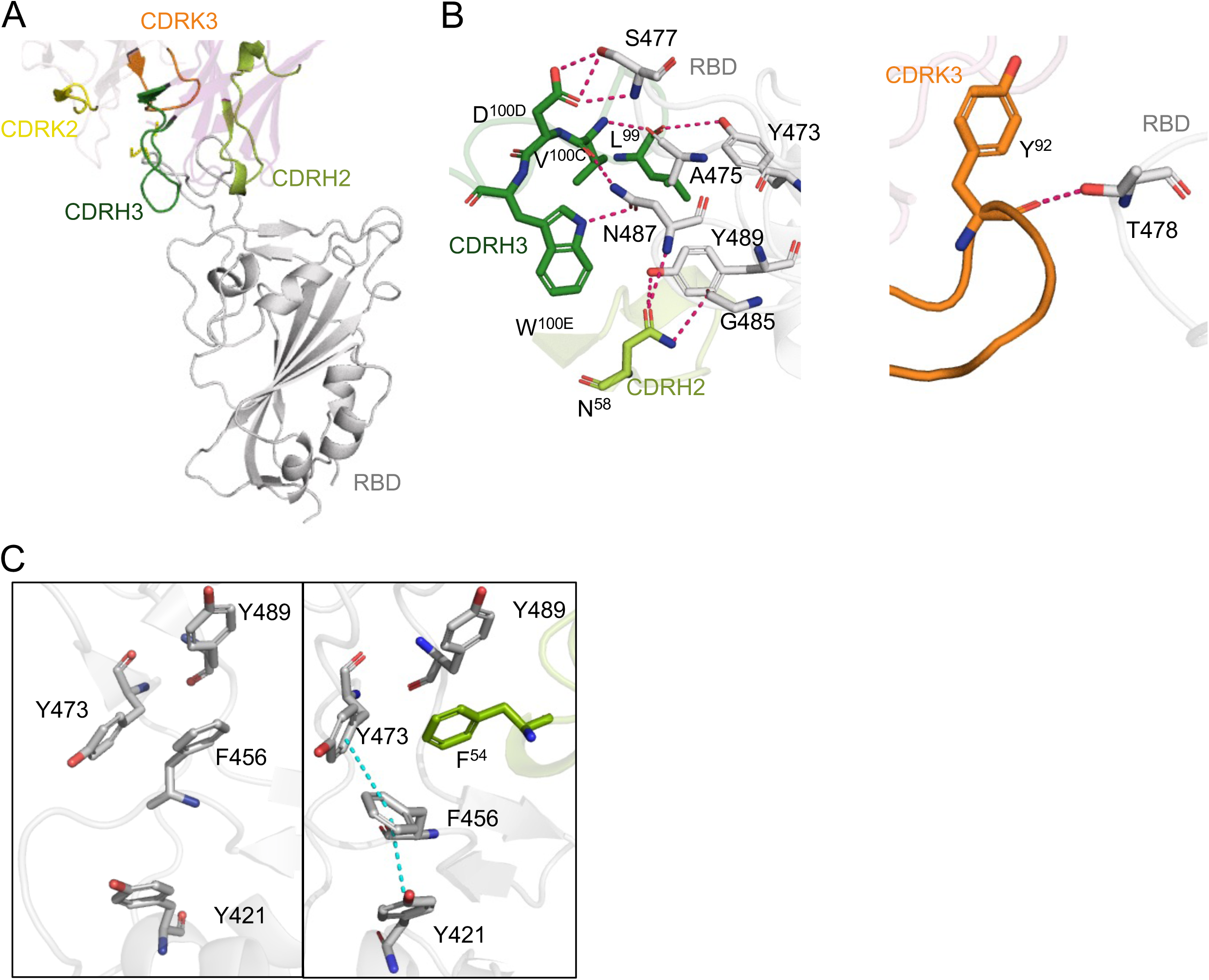
Three-dimensional structure of the 80 Fab-RBD complex. (A) Binding to RBD (grey) involves interactions mediated by heavy chain complementarity determining regions (HCDRs) 2 and 3 (light and dark green, respectively) and kappa light chain CDRS (KCDRs) 2 and 3 (yellow and orange). (B) Detailed view of the hydrogen bond network (dashed lines) formed between key residues at the binding interface of the 80-RBD complex. (C) Rearrangement of RBD aromatic residues (grey) upon 80 binding (green) to form a pi-stacking interaction network (shown in cyan dashed line). Left panel: unbound RBD and right panel: 80-RBD complex.

**Supplementary Figure 8.**
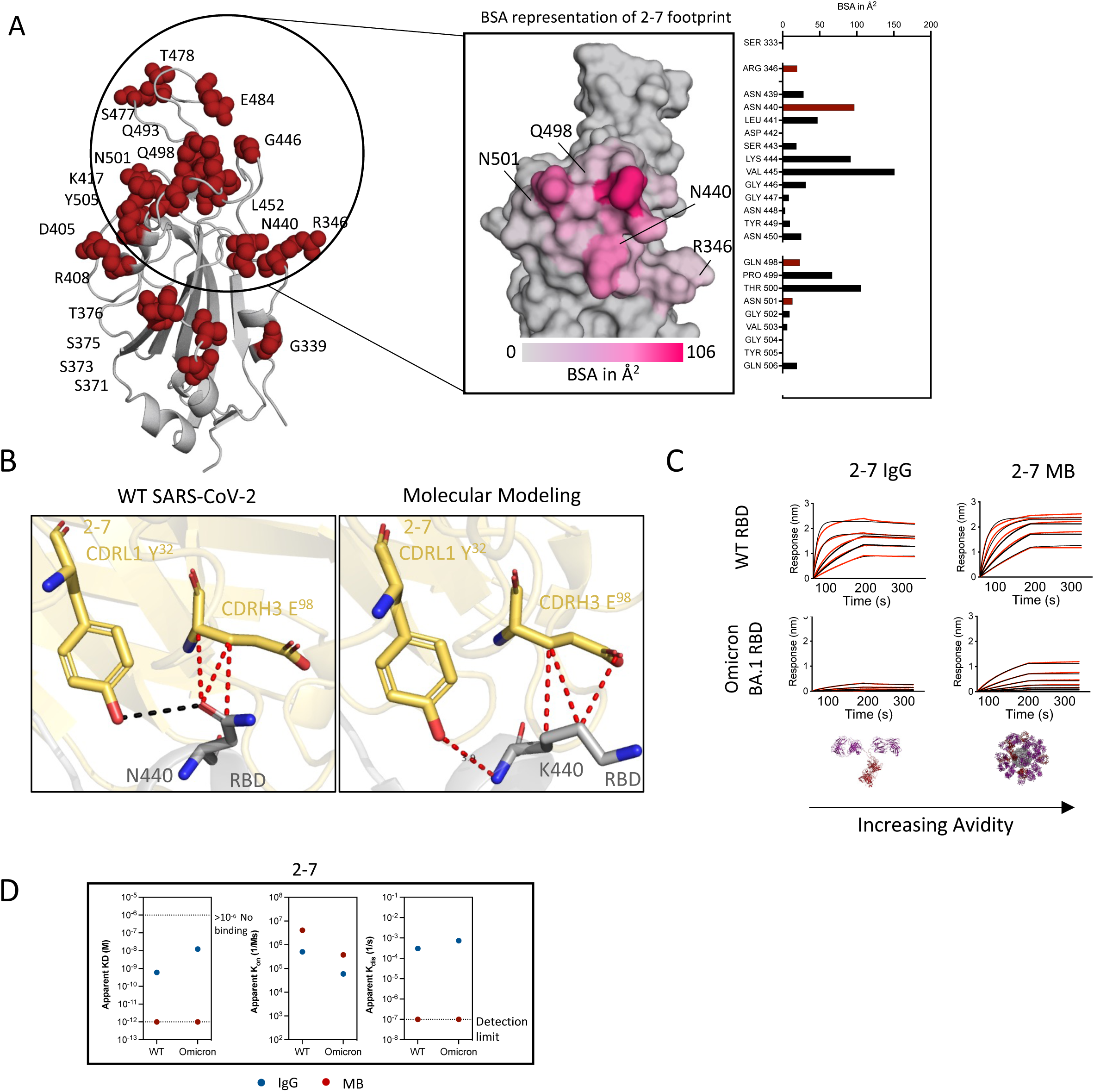
Molecular basis for 2-7 RBD recognition and MB resilience against mutations. (A) Residues commonly mutated in VOCs are represented as red spheres on WT RBD (grey). Inset image shows the surface representation of the RBD with the footprint of 2-7 (PDB ID: 7LSS) colored according to its BSA. Mutated residues in VOC that fall in the antibody footprint are annotated. Right panel - graphical representation of the BSA of residues involved in RBD binding. Residues colored in red are mutated in VOCs. (B) Molecular modeling of the N440K mutation in the RBD-2-7 complex shows that there is sufficient space to accommodate the mutated side chain. Hydrogen bonds and Van der Waals interactions are shown in black and red dashed lines, respectively. (C) Sensograms of 2-7 IgG and MB binding to WT and Omicron BA.1 RBD. Red lines represent raw data and black lines represent global fit. (D) Comparison of the binding kinetic parameters of 2-7 as IgG and MB for binding to WT and Omicron BA.1 RBD. Data shown is average from two independent experiments.

**Supplementary Figure 9.**
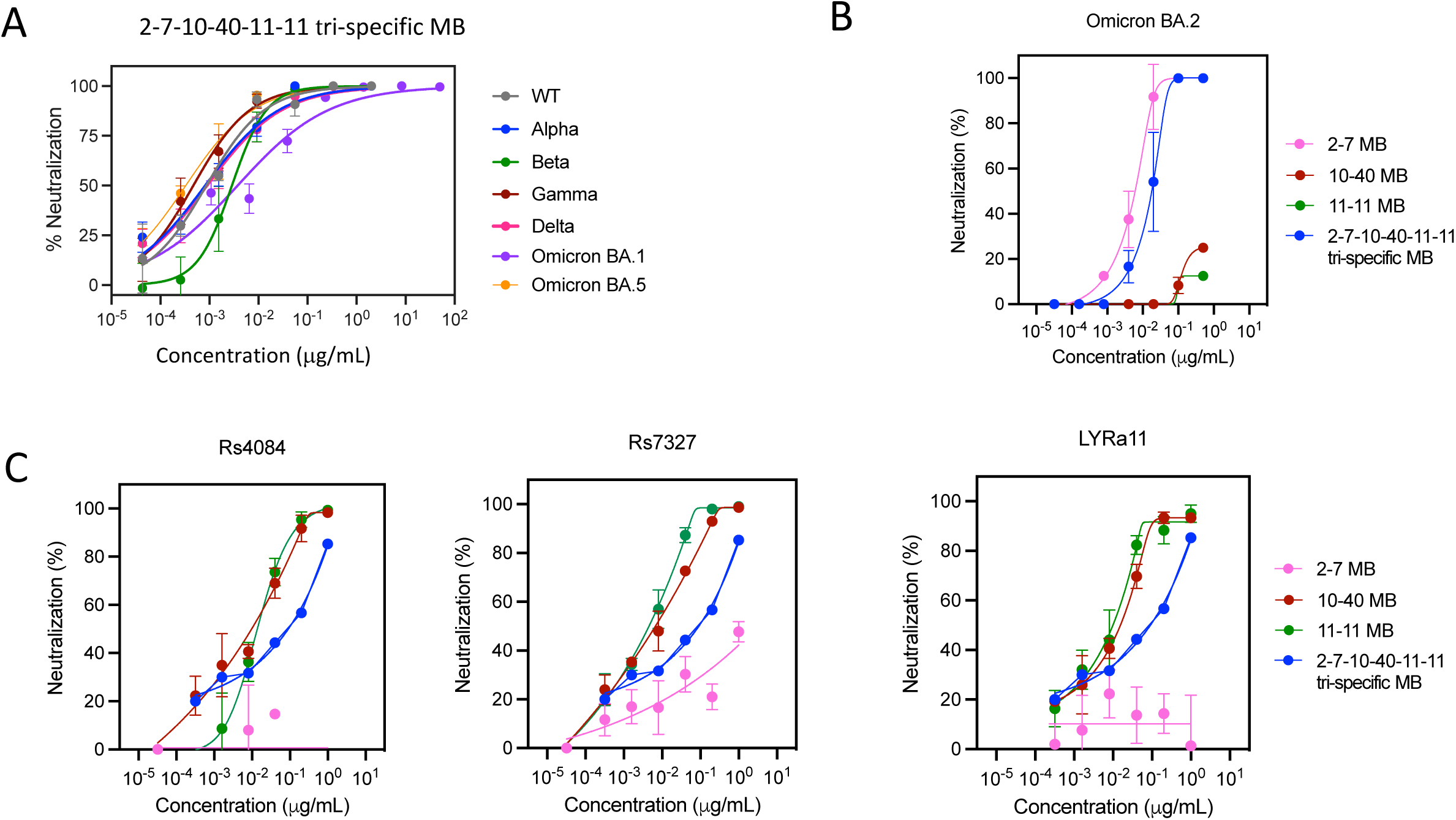
Broad SARS-CoV-2 and sarbecovirus neutralization by tri-specific Multabody. (A) PsV neutralization of 2-7-10-40-11-11 tri-specific MB against SARS-CoV-2 wildtype, Alpha, Beta, Gamma, Delta, Omicron BA.1 and Omicron BA.5 indicated in grey, blue, green, red, pink, purple, and orange, respectively. The mean values ± SEM for three biological replicates are shown in each neutralization plot. (B) Live virus neutralization of 2-7-10-40-11-11 tri-specific MB (blue) and 2-7 (pink), 10-40 (red) and 11-11 (green) mono-specific MBs against Omicron (BA.2) authentic virus. The mean values ± SD for two technical replicates is shown in each neutralization plot. (C) PsV neutralization of 2-7-10-40-11-11 tri-specific MB (blue) and 2-7 (pink), 10-40 (red) and 11-11 (green) mono-specific MBs against Rs4084, Rs7327 and LYRa11 SARS-CoV-1 related bat coronaviruses.

**Supplementary Table 1.**
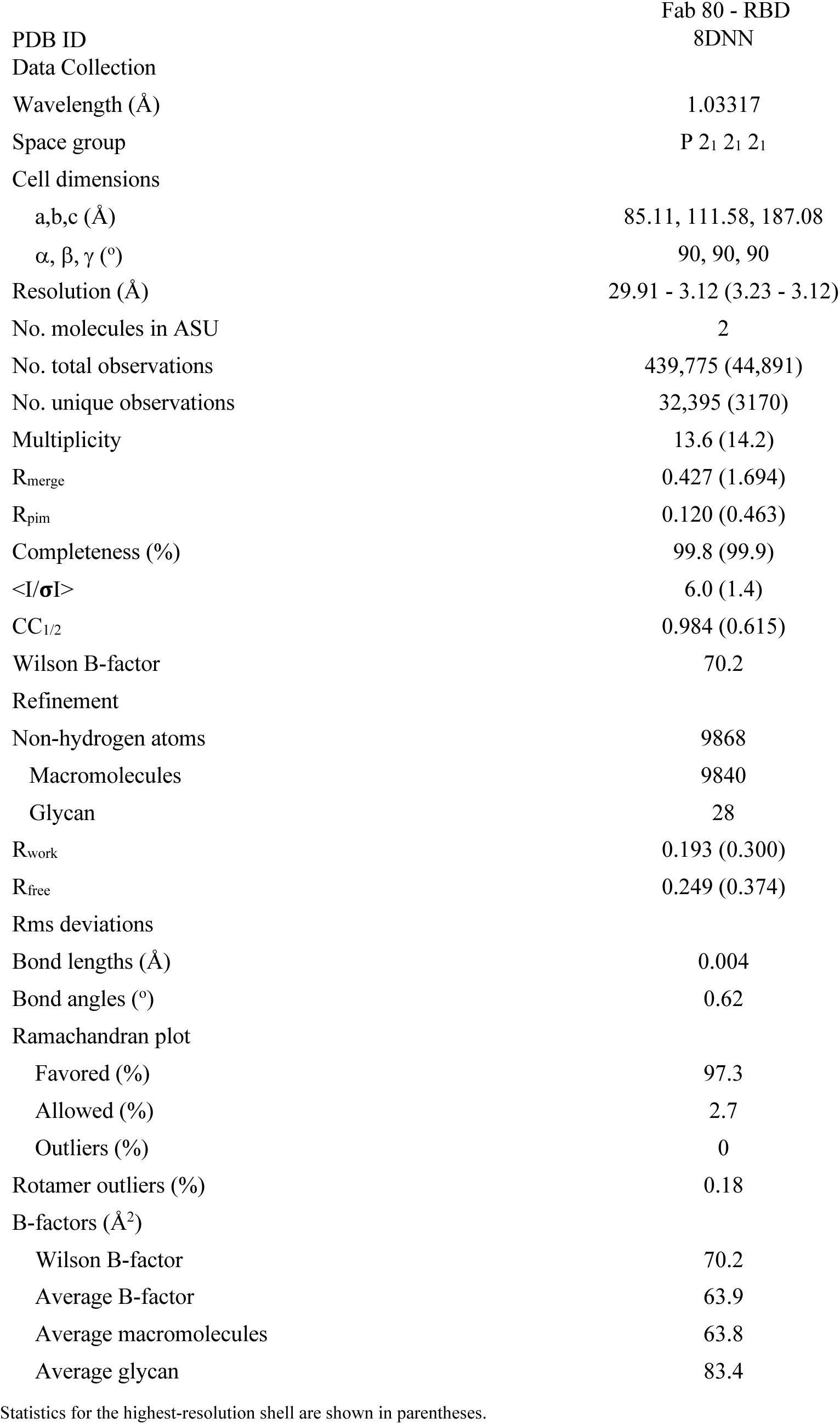
X-ray crystallography data collection and refinement statistics.

|  |  |
| --- | --- |
|  | Fab 80 - RBD |
| PDB ID | 8DNN |
| Data Collection |  |
| Wavelength (Å) | 1.03317 |
| Space group | P 2<sub>1</sub> 2<sub>1</sub> 2<sub>1</sub> |
| Cell dimensions |  |
| a,b,c (Å) | 85.11, 111.58, 187.08 |
| α, β, γ (°) | 90, 90, 90 |
| Resolution (Å) | 29.91 - 3.12 (3.23 - 3.12) |
| No. molecules in ASU | 2 |
| No. total observations | 439,775 (44,891) |
| No. unique observations | 32,395 (3170) |
| Multiplicity | 13.6 (14.2) |
| R<sub>merge</sub> | 0.427 (1.694) |
| R<sub>pim</sub> | 0.120 (0.463) |
| Completeness (%) | 99.8 (99.9) |
| <I/σI> | 6.0 (1.4) |
| CC<sub>1/2</sub> | 0.984 (0.615) |
| Wilson B-factor | 70.2 |
| Refinement |  |
| Non-hydrogen atoms | 9868 |
| Macromolecules | 9840 |
| Glycan | 28 |
| R<sub>work</sub> | 0.193 (0.300) |
| R<sub>free</sub> | 0.249 (0.374) |
| Rms deviations |  |
| Bond lengths (Å) | 0.004 |
| Bond angles (°) | 0.62 |
| Ramachandran plot |  |
| Favored (%) | 97.3 |
| Allowed (%) | 2.7 |
| Outliers (%) | 0 |
| Rotamer outliers (%) | 0.18 |
| B-factors (Å<sup>2</sup>) |  |
| Wilson B-factor | 70.2 |
| Average B-factor | 63.9 |
| Average macromolecules | 63.8 |
| Average glycan | 83.4 |

Statistics for the highest-resolution shell are shown in parentheses.

**Supplementary Table 2.** Fab80-RBD contacting residues identified by PISA.

| RBD | Residue | Interaction | BSA (Å<sup>2</sup>) | Fab 80 (B-HC, C-KC) |
| --- | --- | --- | --- | --- |
| 417 | Lys | vdW | 25 | B-Ile53 |
| 421 | Tyr | vdW | 18 | B-Arg31 |
| 455 | Leu | vdW | 49 | B-Ile53, Phe54 |
| 456 | Phe | vdW | 94 | B-Phe54, Pro100 |
| 473 | Tyr<sup>OH</sup> | HB | 35 | B-Leu99<sup>O</sup> |
|  | Tyr | vdW |  | Pro100 |
| 475 | Ala<sup>O</sup> | HB | 60 | B-VAL100C<sup>N</sup> |
|  | Ala | vdW |  | B-Glu100A |
| 476 | Gly | vdW | 18 | B-Val100B |
| 477 | Ser<sup>N</sup> | HB | 107 | B-Asp100D<sup>OD1</sup> |
|  | Ser<sup>OG</sup> | HB |  | B-Asp100D<sup>OD1</sup> |
|  | Ser<sup>OG</sup> | HB |  | B-Asp100D<sup>OD2</sup> |
|  | Ser | vdW |  | B-Val100B, C-Tyr32 |
| 478 | Thr<sup>OG1</sup> | HB | 47 | C-Tyr92<sup>O</sup> |
| 479 | Pro | vdW | 31 | C-Tyr27D |
| 485 | Gly<sup>O</sup> | HB | 13 | B-Asn58<sup>ND2</sup> |
|  | Gly | vdW |  | C-Ala94 |
| 486 | Phe | vdW | 154 | B-Trp47 |
| 487 | Asn<sup>OD1</sup> | HB | 34 | B-Trp100E<sup>NE1</sup> |
|  | Asn<sup>N</sup> | HB |  | B-Asn58<sup>OD1</sup> |
|  | Asn<sup>ND2</sup> | HB |  | B-Val100C<sup>O</sup> |
|  | Asn | vdW |  | B-Ile52 |
| 489 | Tyr<sup>OH</sup> | HB | 118 | B-Asn58<sup>OD1</sup> |
|  | Tyr | vdW |  | B-Ile52, Phe54, Thr56 |
| 490 | Phe | vdW | 2 | B-Phe54 |

**Supplementary Table 3.** Accession ID for sarbecovirus sequences.

| Sequence ID | Accession | Source |
| --- | --- | --- |
| Rs4084 | KY417144 | NCBI |
| Rs7327 | KY417146 | NCBI |
| LYRa11 | KF569996 | NCBI |

